# ALS-associated RNA binding proteins converge on *UNC13A* transcription through regulation of REST

**DOI:** 10.1101/2024.10.22.619761

**Authors:** Yasuaki Watanabe, Naoki Suzuki, Tadashi Nakagawa, Masaki Hosogane, Tetsuya Akiyama, Hitoshi Warita, Masashi Aoki, Keiko Nakayama

## Abstract

Amyotrophic lateral sclerosis (ALS) is a neurodegenerative disease characterized by the selective loss of motor neurons. Although multiple pathophysiological mechanisms have been identified, a unified molecular basis for ALS has remained elusive. The ALS-associated RNA binding protein (RBP) TDP-43 has previously been shown to stabilize *UNC13A* mRNA by preventing cryptic exon inclusion. We here show that three ALS-associated RBPs—MATR3, FUS, and hnRNPA1—regulate *UNC13A* expression by targeting the silencing transcription factor REST. These three RBPs bind to and downregulate *REST* mRNA and thereby promote *UNC13A* transcription. REST overexpression was detected not only in response to the loss of each of these three RBPs in cultured cells but also in motor neurons of individuals with familial or sporadic ALS. The functional convergence of four RBPs on the regulation of *UNC13A*, a gene essential for synaptic transmission, highlights a pivotal contribution of these proteins to maitainance of synaptic integrity. Our findings thus provide key insight into ALS pathogenesis and a basis for the development of new therapeutic agents.

## Introduction

Amyotrophic lateral sclerosis (ALS) is a rapidly progressive neurodegenerative disease characterized by the degeneration and loss of motor neurons. The identification of several dozen genes that contribute to disease risk or act as causative agents has greatly advanced understanding of ALS pathophysiology (Brown & Al-Chalabi, 2017; Suzuki *et al*, 2023; Watanabe *et al*, 2020). These genes manifest a wide range of predicted functions, and multiple mechanistic hypotheses for their roles in ALS have been proposed, but a unified explanation of ALS pathogenesis has remained elusive. Among the genes mutated in ALS, however, several—including those for TDP-43 (TAR DNA-binding protein–43), MATR3 (matrin 3), FUS (fused in sarcoma), and hnRNPA1 (heterogeneous nuclear ribonucleoprotein A1)—encode RNA binding proteins (RBPs) that share an important role in RNA metabolism (Johnson *et al*, 2014; Kim *et al*, 2013; Van Deerlin *et al*, 2008; Vance *et al*, 2009; Xue *et al*, 2020).

Whereas nuclear loss and cytoplasmic accumulation of TDP-43 are the most prominent features of the pathology of ALS and frontotemporal dementia (Igaz *et al*, 2008; Mitsuzawa *et al*, 2018; Neumann *et al*, 2006), mutations in the COOH-terminal region of FUS result in its mislocalization to the cytoplasm and nuclear depletion, both of which are considered to be central to ALS pathogenesis (Akiyama *et al*, 2019; Suzuki *et al*, 2010; Suzuki *et al*, 2012). Such mislocalization of FUS has been observed in sporadic as well as familial cases of ALS (Tyzack *et al*, 2019). Similarly, the disappearance of nuclear hnRNPA1 and MATR3 has been documented not only in familial ALS but in occasional sporadic cases (Honda *et al*, 2015; Johnson *et al*., 2014; Tada *et al*, 2018). These pathological features suggest that nuclear function of various RBPs is important for the maintenance of motor neurons, and that the loss of such RBP function may lead to convergent impairment of RNA metabolism mediated by these proteins. However, it has remained unclear whether these RBPs regulate a common pathway or target the same RNAs.

In its role as a splicing regulator, TDP-43 suppresses the insertion of cryptic exons during pre-mRNA splicing (Ling *et al*, 2015). If TDP-43 is lost from the nucleus, these cryptic exons remain unspliced, often leading to instability and subsequent degradation of mRNAs via nonsense-mediated decay (NMD). The mRNAs for STMN2 (stathmin 2) and UNC13A (Unc-13 homolog A), both of which are key targets of TDP-43, are especially vulnerable to such instability (Brown *et al*, 2022; Klim *et al*, 2019; Ma *et al*, 2022; Melamed *et al*, 2019). STMN2 dysfunction is associated with both deficient axonal regeneration and the dying-back mechanism of motor neuron death in mice (López-Erauskin *et al*, 2024). UNC13A is essential for the formation and maintenance of synaptic vesicles and neurotransmitter release at synapses (Augustin *et al*, 1999; Willemse *et al*, 2023). Single nucleotide polymorphisms in the noncoding region of *UNC13A* that are associated with increased ALS risk have been found to give rise to the inclusion of a cryptic exon that destabilizes its mRNA (Akiyama *et al*, 2022; van Es *et al*, 2009).

We have now uncovered convergent mechanisms by which four ALS-associated RBPs regulate the expression of *UNC13A*. Whereas TDP-43 stabilizes *UNC13A* mRNA by blocking insertion of a cryptic exon, the loss of MATR3, FUS, or hnRNPA1 gives rise to the transcriptional repression of *UNC13A* mediated by repressor element–1 silencing transcription factor (REST). These findings thus identify *UNC13A* as a key convergence point downstream of these ALS-associated RBPs, suggesting that the dysfunction of these RBPs contributes to a unified pathogenic pathway characterized by loss of UNC13A expression.

## Results

### *UNC13A* expression is downregulated in RBP-KO cell lines

To investigate the possible operation of a common pathway triggered by the dysfunction of RBPs that are genetically and pathologically associated with ALS, we first generated SH-SY5Y human neuroblastoma cell lines deficient in either TDP-43, MATR3, FUS, or hnRNPA1 with the use of the CRISPR/Cas9 system (**Figures S1A–S1D**). RNA–sequencing (seq) analysis of the wild-type (WT) and four RBP–knockout (KO) cell lines revealed transcripts regulated by these RBPs (**Figure 1A**). Application of k-means clustering analysis to the RNA-seq data identified a specific cluster of commonly downregulated genes in the RBP-KO cell lines that was prominently associated with neural processes and synaptic function (**Figure 1B** and **Table S1**), with other clusters being primarily related to nonneuronal processes (**Figure S2**). Further analysis of the cluster of commonly downregulated genes embedded within these neuronal pathways identified *UNC13A*—a gene implicated in the synaptic vesicle cycle—as the only gene previously implicated in ALS (van Es *et al*., 2009; Wroe *et al*, 2008) (**Figure 1C**).

**Figure 1.**
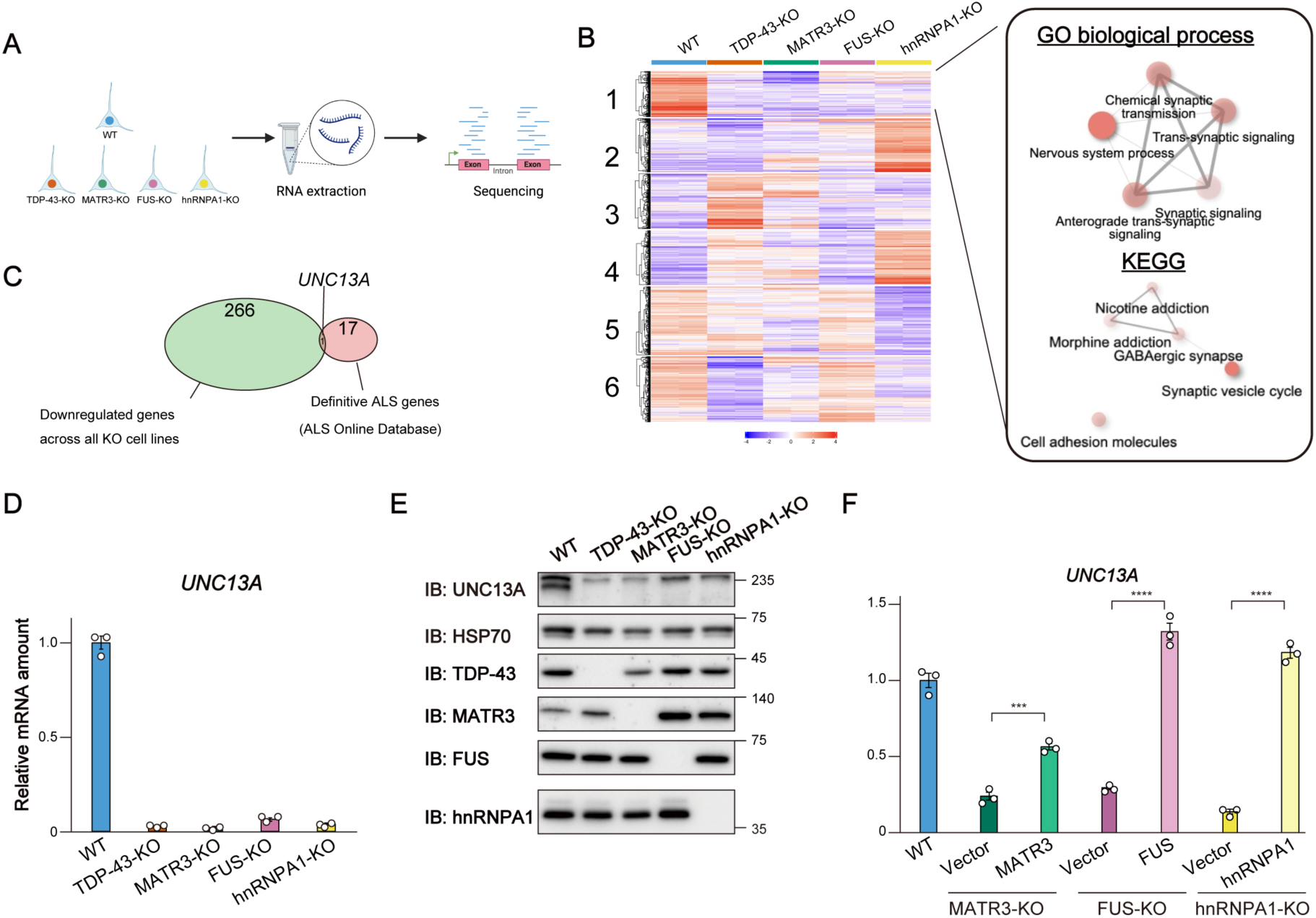
*UNC13A* expression is downregulated in RBP-KO cell lines. (A) Experimental workflow for RNA-seq analysis of SH-SY5Y cell lines deficient in TDP-43, MATR3, FUS, or hnRNPA1. Illustrations were generated with Biorender.com. (B) The results of expression analysis for RNA-seq data from WT as well as TDP-43-, MATR3-, FUS-, and hnRNPA1-KO cell lines. The RNA-seq analysis was performed with biologically independent duplicates. The top 2,000 most differentially expressed genes were classified into six groups by k-means clustering with the use of iDEP version 2.01 (http://bioinformatics.sdstate.edu/idep) (Ge *et al*., 2018), and the data were mean-centered for each gene (left). Genes in cluster 1 were subjected to Gene Ontology (GO) biological process and Kyoto Encyclopedia of Genes and Genomes (KEGG) pathway analysis, and the associated processes and pathways were visualized as a network (right). Color intensity indicates the false discovery rate (FDR) value, representing statistical significance, whereas circle size indicates fold enrichment. (C) Venn diagram showing the overlap between genes in cluster 1 (B) and definitive ALS-related genes with identified mutations in the ALS Online Database (ALSod: http://alsod.iop.kcl.ac.uk) (Wroe *et al*., 2008). (D) RT-qPCR analysis of *UNC13* mRNA in the four RBP-KO cell lines and WT cells. Data are means ± SEM from three independent experiments. (E) Immunoblot (IB) analysis of UNC13A, TDP-43, MATR3, FUS, and hnRNPA1 in the four RBP-KO cell lines and WT cells. HSP70 was examined as a loading control. (F) RT-qPCR analysis of *UNC13A* mRNA in WT cells and in three RBP-KO cell lines complemented with a corresponding doxycycline-inducible RBP vector (or the empty vector as a control) and treated with doxycycline. Data are means ± SEM from three independent experiments. \*\*\**p* < 0.001, \*\*\*\**p* < 0.0001 (Student’s *t* test). See also Figures S1 and S2 and Table S1.

Synaptic dysfunction is thought to contribute to the early stages of ALS (Nishimura & Arias, 2021; Vinsant *et al*, 2013). Given that UNC13A plays a key role in the recycling of synaptic vesicles and neurotransmitter release (Willemse *et al*., 2023), we hypothesized that dysfunction or nuclear depletion of ALS-associated RBPs might lead to a loss of *UNC13A* mRNA and thereby precipitate synaptic dysfunction. To validate our RNA-seq data, we performed reverse transcription and quantitative polymerase chain reaction (RT-qPCR) analysis. This analysis confirmed a substantial reduction in the amount of *UNC13A* mRNA in all four RBP-KO cell lines (**Figure 1D**). In contrast, *STMN2* mRNA, another key target of TDP-43, was depleted only in TDP-43-KO cells (**Figure S1E**). Immunoblot analysis revealed that the UNC13A protein was also depleted in MATR3-, FUS-, and hnRNPA1-KO cell lines as well as in TDP-43-KO cells (**Figure 1E**). The reintroduction of the respective RBP cDNA into MATR3-, FUS-, or hnRNPA1-KO cell lines restored *UNC13A* mRNA abundance (**Figures 1F** and **S1F–S1H**), indicating that the loss of *UNC13A* mRNA was not due to off-target effects of each RBP gene knockout. Collectively, these findings suggested that ALS-associated RBPs commonly regulate synapse-related gene expression including that of *UNC13A*.

### TDP-43 stabilizes *UNC13A* mRNA by blocking cryptic splicing, whereas MATR3, FUS, and hnRNPA1 promote *UNC13A* transcription

Nuclear depletion of TDP-43 has been associated with the downregulation of *UNC13A* expression attributable to cryptic exon inclusion and consequent mRNA instability (Brown *et al*., 2022; Ma *et al*., 2022). To investigate whether the loss of *UNC13A* mRNA observed in cells depleted of other ALS-associated RBPs was also dependent on cryptic exon inclusion, we performed RT-qPCR analysis to detect the cryptic exon included in mature *UNC13A* mRNA in response to TDP-43 depletion (Ma *et al*., 2022). With the use of primers designed specifically for the detection of this exon, we found it to be present in *UNC13A* transcripts of TDP-43-KO cells, but not in those of MATR3-, FUS-, or hnRNPA1-KO cells (**Figure 2A**). This observation indicated that the downregulation of *UNC13A* expression apparent in these latter cell lines is not dependent on cryptic exon insertion.

**Figure 2.**
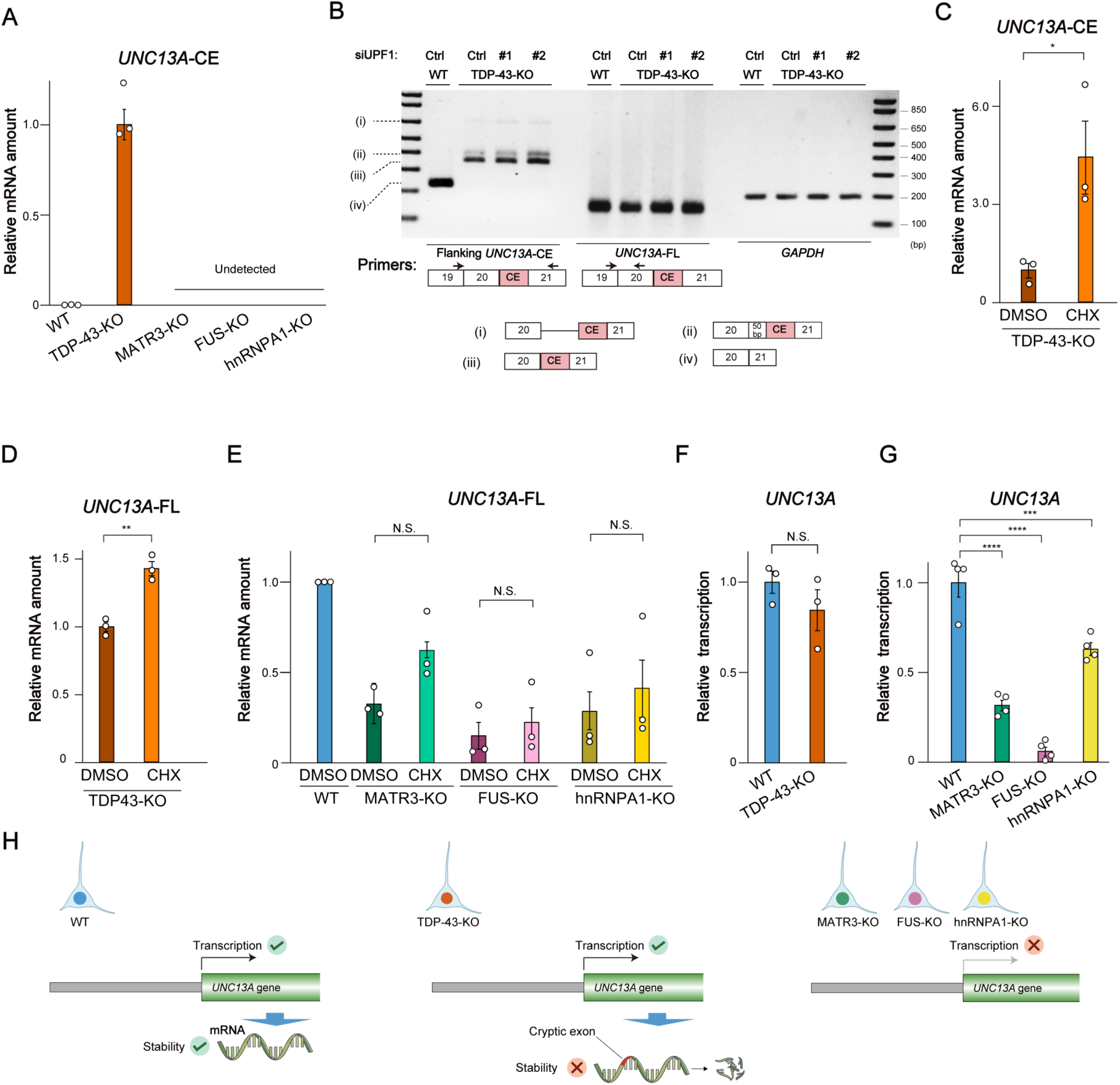
TDP-43 stabilizes *UNC13A* mRNA by blocking cryptic splicing, whereas MATR3, FUS, and hnRNPA1 promote *UNC13A* transcription. (A) RT-qPCR analysis of *UNC13A* transcripts including the cryptic exon (CE) in WT cells and the four RBP-KO cell lines. Data are means ± SEM from three independent experiments. (B) RT-PCR analysis of WT or TDP-43-KO cells transfected with either a GC duplex (negative control, Ctrl) or two different small interfering RNAs (siRNAs #1 or #2) for *UPF1*. The PCR primer locations are shown below the gel images; they flanked the CE of *UNC13A* or recognized a region of *UNC13A* mRNA unaffected by CE inclusion (FL). *GAPDH* was examined as an internal control. Among the PCR products amplified with the primers flanking the CE of *UNC13A*, (i) to (iii) indicate different intron retention patterns for products containing the CE, whereas (iv) indicates a product lacking the CE. (C–E) RT-qPCR analysis of *UNC13A*-CE mRNA in TDP-43-KO cells (C), *UNC13A*-FL mRNA in TDP-43-KO cells (D), and *UNC13A*-FL mRNA in WT, MATR3-KO, FUS-KO, and hnRNPA1-KO cells (E) after treatment with either cycloheximide (CHX, 100 µg/ml) or dimethyl sulfoxide (DMSO) vehicle for 6 h. Data are means ± SEM from three independent experiments. \**p* < 0.05, \*\**p* < 0.01; N.S., not significant (Student’s *t* test). (F and G) RT-qPCR analysis of nascent *UNC13A*-FL mRNA in WT and TDP-43-KO cells (F) as well as in WT, MATR3-KO, FUS-KO, and hnRNPA1-KO cells (G) that had been labeled with 4-EU. Data are means ± SEM from three (F) or four (G) independent experiments. \*\*\**p* < 0.001, \*\*\*\**p* < 0.0001, N.S. (Student’s *t* test). (H) Proposed mechanisms for the regulation of *UNC13A* mRNA abundance in WT, TDP-43-KO, and the other three types of RBP-KO cells. Illustrations were generated with Biorender.com. See also **Figure S3**.

To confirm that *UNC13A* mRNA containing the cryptic exon is degraded via the NMD pathway in our TDP-43-KO cell model, we examined the effect of NMD inhibition on *UNC13A* mRNA stability. RT-PCR analysis with primers designed to amplify regions including the cryptic exon revealed the presence of PCR products containing this exon in TDP-43-KO cells (**Figure 2B**). The amount of these PCR products was increased by knockdown of UPF1 (**Figures 2B** and **S3A**), a key component of the NMD pathway. Furthermore, treatment of TDP-43-KO cells with the NMD inhibitor cycloheximide restored *UNC13A* mRNA levels, as confirmed by RT-qPCR analysis targeting both cryptic and canonical exons (**Figures 2C, 2D**, and **S3B**). These findings confirmed that *UNC13A* mRNA is degraded as a result of the inclusion of a cryptic exon and subsequent NMD in the absence of TDP-43.

In contrast, inhibition of NMD in MATR3-, FUS-, or hnRNPA1-KO cells did not significantly affect *UNC13A* mRNA abundance (**Figures 2E** and **S3C**). To compare nascent *UNC13A* transcript levels between the four RBP-KO cell lines and WT cells, we exposed the cells to 4-thiouridine (4-EU) to allow its incorporation into the newly synthesized transcriptome. Whereas 4-EU–labeled nascent *UNC13A* transcript levels were similar in TDP-43-KO cells and WT cells, they were significantly reduced in the other three RBP-KO cell lines (**Figures 2F** and **2G**). Further RT-qPCR analysis with primers targeting intronic regions of *UNC13A* did not show a reduction in the amount of *UNC13A* pre-mRNA in TDP-43-KO cells compared with WT cells, but a significant reduction was observed in MATR3-, FUS-, and hnRNPA1-KO cell lines (**Figures S3D** and **S3E**). These findings indicated that, under normal physiological conditions, ALS-associated RBPs regulate *UNC13A* mRNA abundance through distinct mechanisms. Specifically, in the absence of TDP-43, *UNC13A* mRNA is destabilized as a result of the inclusion of a cryptic exon and degraded via the NMD pathway. In contrast, in the absence of MATR3, FUS, or hnRNPA1, *UNC13A* transcription is disrupted (**Figure 2H**).

### REST is upregulated and binds to the *UNC13A* promoter in RBP-KO cells

The dysregulation of *UNC13A* transcription in MATR3-, FUS-, and hnRNPA1-KO cells did not likely reflect a direct effect of these proteins on gene transcription given their established roles as RBPs (Xue *et al*., 2020). We therefore hypothesized that these RBPs influence a specific transcription factor that regulates *UNC13A* mRNA synthesis. To identify such a transcription factor, we performed a meta-analysis of chromatin immunoprecipitation (ChIP)–seq data sets from the ENCODE database (2012; Oki *et al*, 2018). Sixteen data sets showed significant peaks indicative of transcription factor binding at the *UNC13A* promoter region, with five proteins being enriched in this region (**Figure 3A** and **Table S2**). Among these five proteins, REST has been shown to suppress neuronal gene expression in nonneural tissues, a key function required for proper activation of these genes only in appropriate cells (Andrés *et al*, 1999; Lunyak & Rosenfeld, 2005). Disruption of this REST-mediated regulatory mechanism has been implicated in neurodegenerative disease (Andrés *et al*., 1999; Soldati *et al*, 2013). The ChIP-seq data revealed that REST binds to the *UNC13A* promoter in both neuroblastoma and nonneuronal cell lines, and we confirmed this binding pattern in SH-SY5Y cells by ChIP-qPCR analysis (**Figures S4A** and **S4B**).

**Figure 3.**
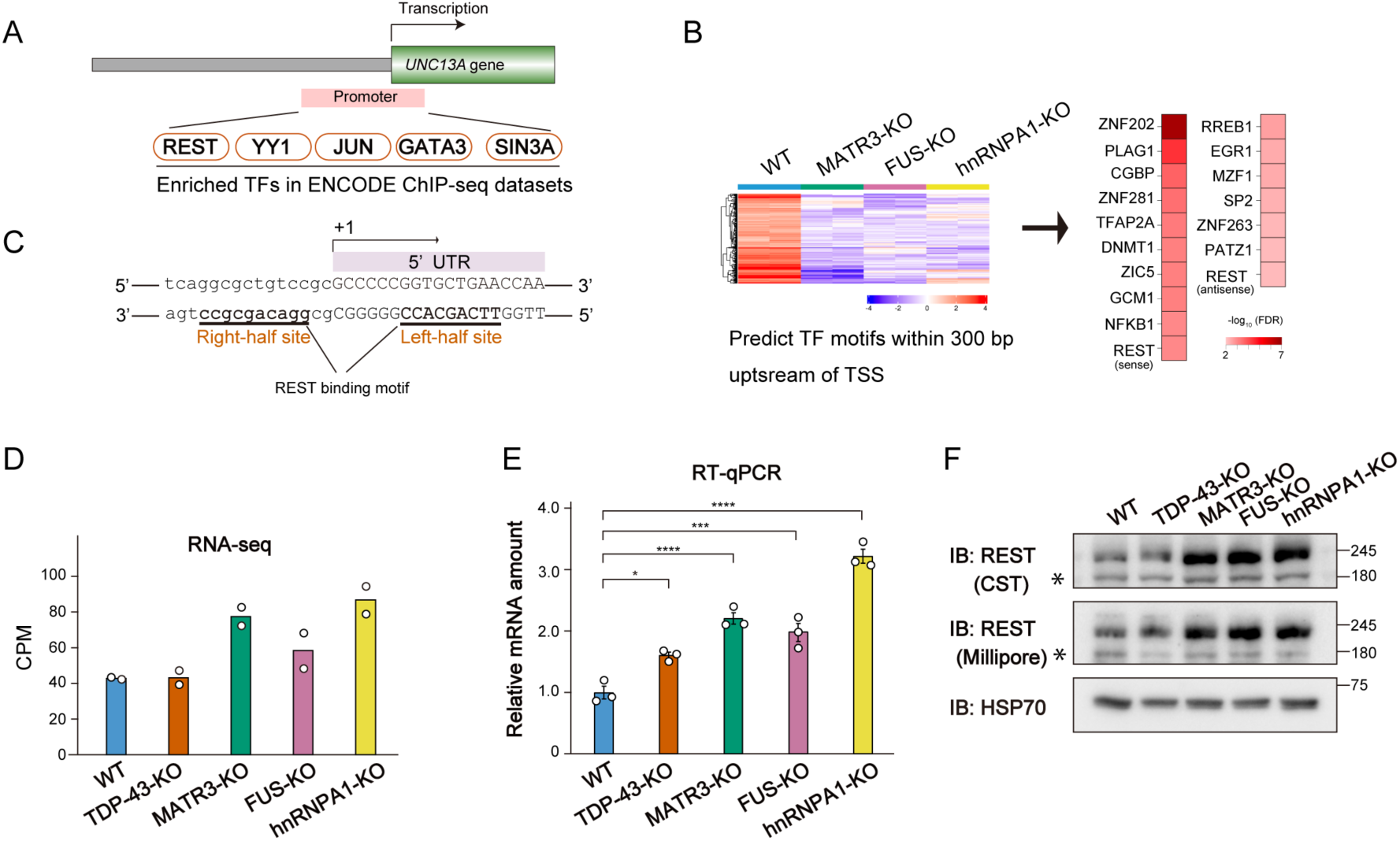
REST is upregulated and binds to the *UNC13A* promoter in RBP-KO cells. (A) Identification of transcription factors (TFs) that bind to the *UNC13A* promoter with the use of ChIP-Atlas (Oki *et al*., 2018). The *UNC13A* promoter region was aligned with ChIP-seq data sets from ENCODE that are specific to neuronal cells in order to identify enriched transcription factors. The enrichment score threshold was set at 500, and the analyzed promoter region was a 339-bp ENCODE candidate cis-regulatory element (cCRE) corresponding to chr19:17688234-17688572 in Hg38. The five identified transcriptional factors are shown. UTR, untranslated region. (B) Motif analysis for transcription factors enriched in the promoter regions (300 bp upstream of the TSS) of genes commonly downregulated in MATR3-, FUS-, and hnRNPA1-KO cell lines. The downregulated genes were identified by k-means clustering (left). Transcription factors with an FDR of <0.01 are shown in a heat map based on the FDR values (right). (C) REST binding motif within the *UNC13A* promoter region. Bases corresponding to the noncanonical motif on the antisense strand are underlined. (D and E) *REST* mRNA abundance in four RBP-KO cell lines and WT cells as determined by RNA-seq (D) or RT-qPCR (E) analysis. The RNA-seq data are means from biological duplicates and are presented as counts per million (CPM), and the RT-qPCR data are means ± SEM from three independent experiments. \**p* < 0.05, \*\*\**p* < 0.001, \*\*\*\**p* < 0.0001 (one-way ANOVA followed by Tukey’s post hoc test). (F) Immunoblot analysis of REST in four RBP-KO cell lines and in WT cells. The REST protein was detected at a position corresponding to ∼210 kDa with two different antibody preparations. Asterisks indicate nonspecific bands. Quantitative data are presented in **Figure S4C**. See also **Figure S4** and **Tables S2** to **S4**.

For subsequent analysis of RNA-seq data, we focused on genes with transcript levels downregulated as for *UNC13A* in MATR3-, FUS-, and hnRNPA1-KO cells in order to search for specific transcriptional regulators in these three cell lines. We identified common transcription factor binding motifs within 300 bp of the transcription start site (TSS) of such genes. These motifs included two complementary sequences targeted by REST (**Figure 3B** and **Table S3**), further emphasizing the extensive regulatory influence of this transcription factor. In addition, we noticed the presence of a noncanonical NRSE motif for REST binding in the *UNC13A* promoter region (Johnson *et al*, 2008) (**Figure 3C**).

On the basis of these findings, we hypothesized that REST might be upregulated in RBP-KO cells compared with WT cells and thereby repress transcription of *UNC13A* and other target genes. Indeed, RNA-seq and RT-qPCR analyses confirmed that *REST* mRNA abundance was substantially higher in MATR3-, FUS-, and hnRNPA1-KO cells than in WT cells (**Figures 3D** and **3E**). In addition, the amount of REST protein was increased in these three RBP-KO cell lines but not in TDP-43-KO cells (**Figures 3F** and **4C**), suggesting that upregulation of REST might play a key role in the control of *UNC13A* and other gene expression in the absence of MATR3, FUS, or hnRNPA1.

We also investigated whether genes whose transcription is regulated by REST are included among the commonly downregulated genes in MATR3-, FUS-, and hnRNPA1-KO cells. We defined potential REST target genes as genes with a REST binding site within ±1 kb of the TSS by cross-referencing ENCODE data sets (2012; Oki *et al*., 2018). We identified 38 genes that were both potential targets of REST and downregulated in the three RBP-KO cell lines (**Figure S4D** and **Table S4**). GO analysis of these genes identified several including *UNC13A* as being associated with synapse-related functions (**Figure S4E**), suggesting that dysfunction of MATR3, FUS, and hnRNPA1 commonly induces synaptic pathology as a result of dysregulation of REST.

### REST inhibits *UNC13A* transcription in RBP-KO cells

To assess the impact of REST upregulation on *UNC13A* promoter activity, we performed a luciferase reporter assay with three different constructs: a promoter-less luciferase (mock) construct as a control, a canonical construct containing the entire *UNC13A* promoter region, and a modified (ΔR) construct that contains a version of the *UNC13A* promoter lacking a 6-bp sequence that is required for REST binding and possesses a high PhyloP score indicative of a high level of evolutionary conservation (**Figures 4A** and **4B**). The promoter activity of the ΔR construct was significantly increased compared with that of the WT (canonical) construct in HEK293T cells, suggesting that endogenous REST represses *UNC13A* transcription (**Figure 4C**). Furthermore, whereas knockdown of REST mediated by the CRISPR/Cas9 system increased the activity of the canonical construct, it had no significant effect on that of the ΔR construct (**Figures 4D** and **4E**). These data suggested that REST represses *UNC13A* transcription through its interaction with the gene promoter.

**Figure 4.**
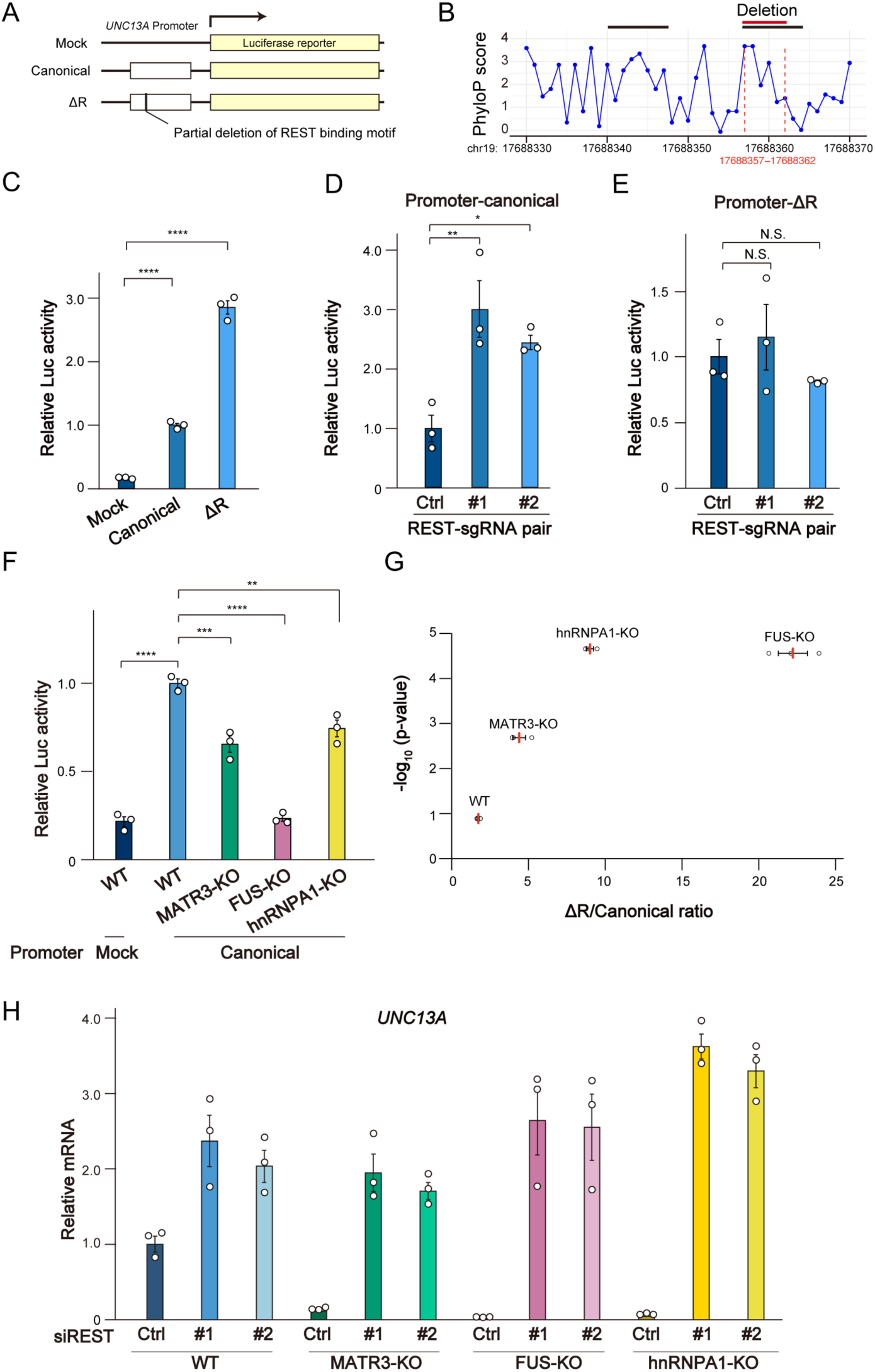
REST inhibits *UNC13A* transcription in RBP-KO cells. (A) Schematic diagram of mock, canonical, and ΔR luciferase reporter constructs for the *UNC13A* promoter. The ΔR construct lacks a 6-bp sequence essential for REST binding. (B) Line graph showing PhyloP scores for evolutionary conservation of the *UNC13A* promoter region. The black bars indicate the REST binding motif, and the red bar indicates the region deleted in the ΔR construct. (C) Relative luciferase (Luc) activity for HEK293T cells transfected with the firefly luciferase constructs shown in (A) as well as with a vector for *Renilla* luciferase. The firefly/*Renilla* luciferase activity ratio was measured 2 days after transfection. Data are means ± SEM from three independent experiments. \*\*\*\**p* < 0.0001 (one-way ANOVA followed by Tukey’s post hoc test). (D and E) Relative luciferase activity for the canonical (D) and ΔR (E) promoter constructs in HEK293T cells transfected previously with two different pairs of Cas9–single guide RNA (sgRNA) vectors targeting REST or with a control vector (Ctrl). Data are means ± SEM from three independent experiments. \**p* < 0.05, \*\**p* < 0.01, NS (one-way ANOVA followed by Tukey’s post hoc test). (F) Relative luciferase activity for the mock and canonical promoter constructs in WT and RBP-KO cell lines. Data are means ± SEM from three independent experiments. \*\**p* < 0.01, \*\*\**p* < 0.001, \*\*\*\**p* < 0.0001 (one-way ANOVA followed by Tukey’s post hoc test). (G) The ΔR/canonical promoter construct ratio of luciferase activity in WT and RBP-KO cell lines (*x*-axis). The vertical axis shows the –log_10_(*p* value) for comparison between the activities of the canonical and ΔR constructs in each cell line (Student’s *t* test). Ratio data are means (red bars) ± SEM from three independent experiments. (H) RT-qPCR analysis of *UNC13A* mRNA in WT, MATR3-KO, FUS-KO, and hnRNPA1-KO cell lines transfected with a GC duplex (negative control) or two different siRNAs targeting *REST*. Data are means ± SEM from three independent experiments. See also **Figure S5**.

The activity of the canonical construct was significantly lower in MATR3-, FUS-, and hnRNPA1-KO cell lines compared with WT cells (**Figure 4F**), consistent with the observation that *UNC13A* transcription is downregulated in these three RBP-KO cell lines (**Figures 2G** and **S3E**). Conversely, the activity of the ΔR construct was increased in the three RBP-KO cell lines, with the ΔR/canonical activity ratio far exceeding that for WT cells (**Figure 4G**), suggesting that the inability of REST to bind to the mutated *UNC13A* promoter results in a large increase in *UNC13A* transcriptional activity in MATR3-, FUS-, and hnRNPA1-KO cells, in which REST is overexpressed relative to WT cells.

To confirm that this regulation of *UNC13A* transcription by REST is reflected in the amount of *UNC13A* mRNA in WT and RBP-KO cell lines, we depleted *REST* mRNA in the cells by ∼90% by siRNA transfection (**Figure S5A**). This intervention resulted in a significant increase in *UNC13A* expression, which achieved similar levels in all tested cell lines (**Figure 4H**). Together, these data suggested that the downregulation of *UNC13A* transcription apparent in MATR3-, FUS-, and hnRNPA1-KO cell lines is attributable to REST overexpression. In contrast, such REST knockdown had no significant effect on *UNC13A* expression in TDP-43-KO cells (**Figures S5B** and **S5C**), consistent with our observation that *UNC13A* mRNA is destabilized and degraded as a result of cryptic exon insertion in these cells (**Figures 2A–2D**).

### MATR3, FUS, and hnRNPA1 bind to *REST* mRNA

On the basis of our finding that *REST* mRNA abundance is increased in MATR3-, FUS-, and hnRNPA1-KO cell lines (**Figures 3D** and **3E**), we investigated the mechanism by which these three RBPs regulate *REST* expression. We found that the level of nascent *REST* mRNA was unchanged in the three RBP-KO cell lines compared with WT cells (**Figure 5A**), suggesting that the increase in the amount of *REST* mRNA in the RBP-KO cells was not due to an effect on transcription. We next performed an RNA immunoprecipitation (RIP) assay to determine whether the three RBPs interact with *REST* mRNA. This analysis showed that MATR3, FUS, and hnRNPA1 indeed bound to *REST* mRNA (**Figures 5B–5E**), suggesting that the regulation of *REST* mRNA abundance by these RBPs occurs at the posttranscriptional level through binding to and consequent destabilization of *REST* mRNA.

**Figure 5.**
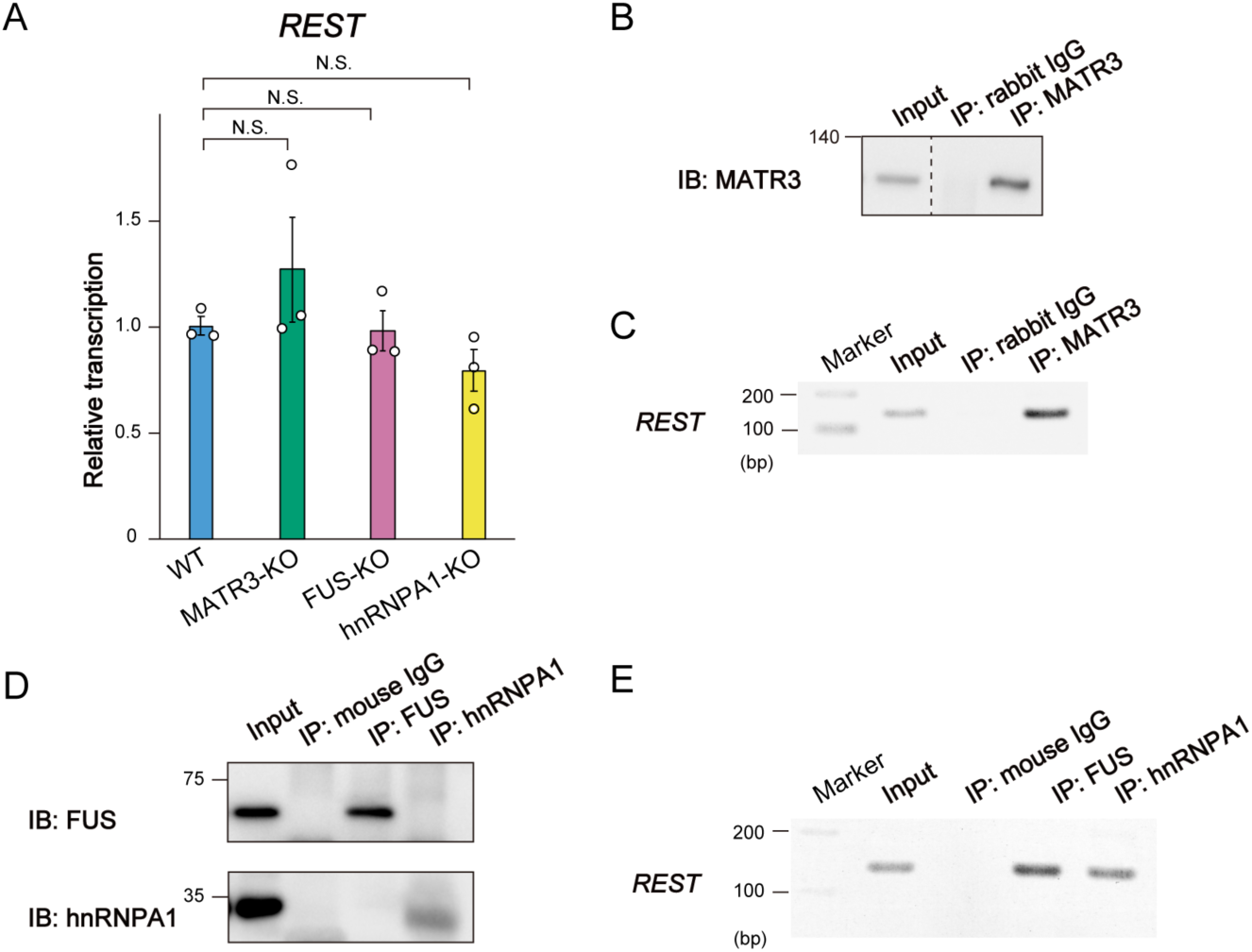
MATR3, FUS, and hnRNPA1 bind to *REST* mRNA. (A) RT-qPCR analysis of nascent *REST* mRNA abundance in WT, MATR3-KO, FUS-KO, and hnRNPA1-KO cell lines. Data are means ± SEM from three independent experiments. N.S. (one-way ANOVA followed by Tukey’s post hoc test). (B) Enrichment of MATR3 by immunoprecipitation from WT cell lysate. The cell lysate (Input) as well as the immunoprecipitate (IP) obtained with antibodies to MATR3 or with control immunoglobulin G (IgG) were subjected to immunoblot analysis with antibodies to MATR3. (C) Detection of *REST* mRNA by RT-PCR analysis of the samples obtained as in (B). (D) Enrichment of FUS and hnRNPA1 by immunoprecipitation from WT cell lysate. (E) Detection of *REST* mRNA by RT-PCR analysis of the samples obtained as in (D).

### FUS phase separation underlies regulation of *REST* mRNA

Among the three RBPs that influence REST expression, we next focused on FUS in order to further investigate the molecular mechanism of *REST* mRNA regulation, given that depletion of FUS attenuated transcriptional regulation of *UNC13A* to a greater extent than did that of the other RBPs (**Figures 2G, 4F**, and **S3E**) and that *FUS* is more frequently mutated in individuals with ALS compared with the other RBP genes (Akiyama *et al*, 2016; Renton *et al*, 2014).

To identify which portions of the RNA binding domain (amino acids 212–526) of FUS influence *REST* mRNA stability, we generated expression vectors for full-length (FL) FUS and deletion mutants lacking the glycine-rich, RRM, RGG1, RGG2, or ZnF domains (ΔGly, ΔRRM, ΔRGG1, ΔRGG2, and ΔZnF, respectively) (**Figure 6A**) and introduced these constructs into FUS-KO cells (**Figure 6B**). Although the expression level of FL in FUS-KO cells was relatively low compared with that of the endogenous protein in WT cells, its expression in FUS-KO cells effectively reduced the abundance of *REST* mRNA to a level similar to that apparent in WT cells (**Figures 6B** and **6C**). Whereas, like FL, the ΔRRM and ΔZnF mutants suppressed the amount of *REST* mRNA and rescued *UNC13A* expression in FUS-KO cells, the ΔGly, ΔRGG1, and ΔRGG2 mutants had no such effects (**Figures 6D** and **6E**). The domains deleted in the latter three mutants showed high PONDR scores, suggestive of the presence of intrinsically disordered regions (IDRs) (**Figure 6A**). Given that IDRs contribute to phase separation,(Molliex *et al*, 2015) the inability of these three mutants lacking IDRs to compensate for the loss of endogenous FUS implicated phase separation in the regulation of *REST* mRNA by FUS.

**Figure 6.**
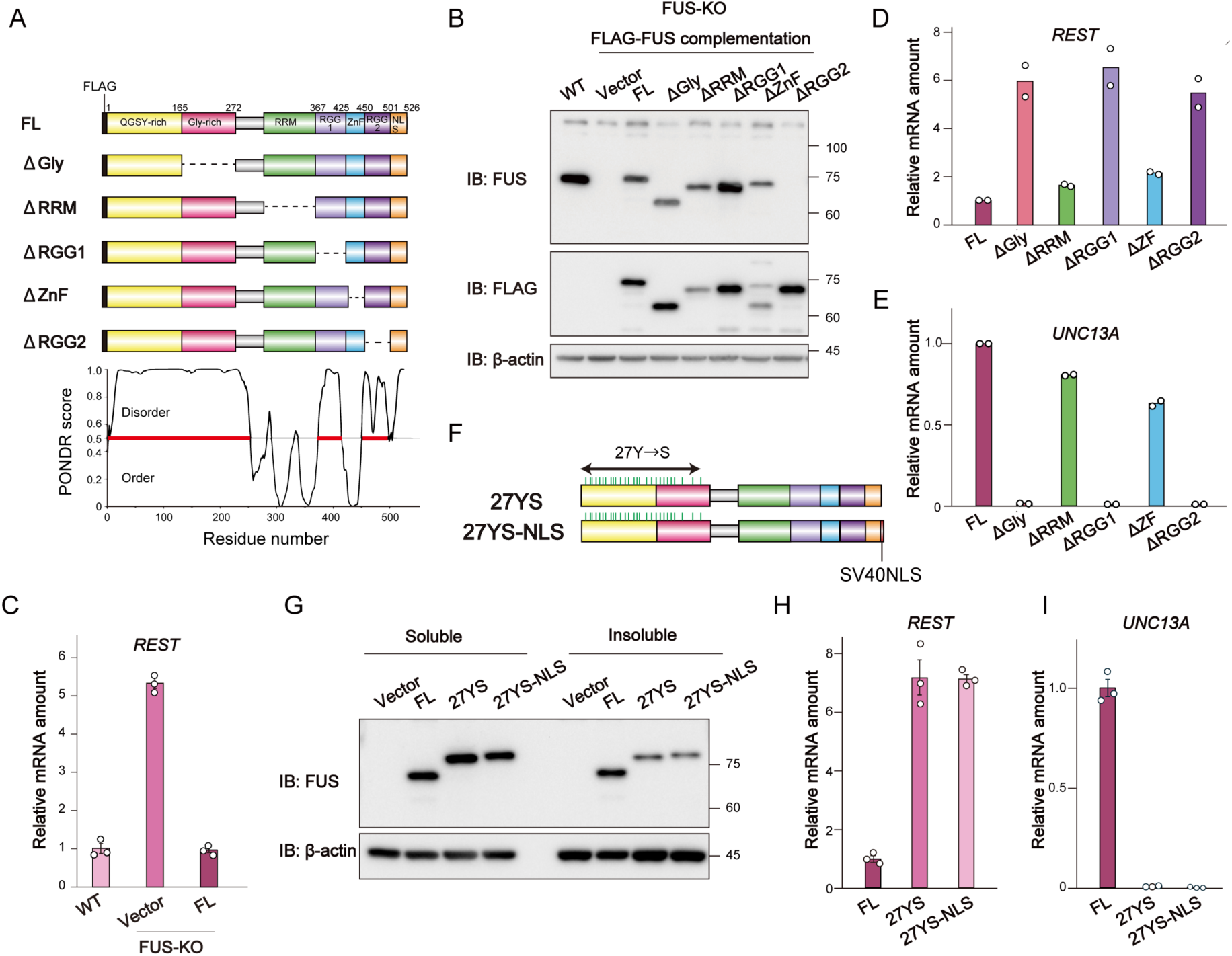
FUS phase separation underlies regulation of *REST* mRNA. (A) Schematic representation of FLAG epitope–tagged human FUS mutant constructs (top). Dashed lines indicate deleted domains. FL, full length; Gly, glycine-rich domain; RRM, RNA recognition motif; RGG, Arg-Gly-Gly; ZnF, zinc finger. Disorder prediction for FUS residues by PONDR (http://www.pondr.com) is shown at the bottom. (B) Immunoblot analysis of endogenous FUS in WT cells and of ectopic FLAG-tagged FL or deletion mutant forms of FUS expressed in FUS-KO cells. β-actin was examined as a loading control. (C) RT-qPCR analysis of *REST* mRNA in WT cells as well as in FUS-KO cells expressing FUS-FL or harboring the empty vector. Data are means ± SEM from three independent experiments. (D and E) RT-qPCR analysis of *REST* (D) and *UNC13A* (E) mRNAs in FUS-KO cells expressing FL or deletion mutant forms of FUS. Data are means from two independent experiments. (F) Schematic representation of FUS mutants deficient in phase separation activity. The 27 NH_2_-terminal tyrosines are all replaced by serine in 27YS, whereas 27YS-NLS also possesses the NLS of SV40 at its COOH-terminus. (G) Immunoblot analysis of FUS in FUS-KO cells expressing FL, 27YS, or 27YS-NLS forms of FUS. The cell lysates were prepared in the presence of 0.5% Nonidet P-40 detergent and centrifuged at 20,000 × *g* for 15 min at 4°C, and the resulting supernatant was collected as the soluble fraction. The pellet was subjected to ultrasonic treatment and dissolved in urea buffer to obtain the insoluble fraction. (H and I) RT-qPCR analysis of *REST* (H) and *UNC13A* (I) mRNAs in FUS-KO cells expressing FL, 27YS, or 27YS-NLS forms of FUS. Data are means ± SEM from three independent experiments.

The 27 tyrosine residues in the NH_2_-terminal prionlike domain (PrLD, amino acids 1–239) of FUS mediate phase separation through interaction with arginine residues in the IDRs of the RNA binding domain (Qamar *et al*, 2018; Wang *et al*, 2018). To investigate the role of FUS phase separation in the regulation of *REST* mRNA, we substituted these 27 tyrosine residues with serine (27YS mutant) to impair such separation (**Figure 6F**). To prevent loss of the 27YS mutant from the nucleus as a result of an inability to interact with chromatin (Reber *et al*, 2021), we also incorporated the SV40 nuclear localization signal into the mutant protein (27YS-NLS mutant). Expression of the 27YS and 27YS-NLS mutants in FUS-KO cells revealed that, unlike the FL protein, they were predominantly found in the soluble fraction of cell lysates (**Figure 6G**), indicating that phase separation is required for the formation of insoluble aggregates. Neither mutant attenuated *REST* expression or restored *UNC13A* expression in the FUS-KO cells (**Figures 6H** and **6I**). These results thus underscored the essential role of phase separation in the regulation of *REST* mRNA stability by FUS.

### REST is overexpressed in spinal motor neurons of individuals with ALS

Our data showed that *UNC13A* expression is repressed due to the overexpression of *REST* in SH-SY5Y cells depleted of MATR3, FUS, or hnRNPA1. To validate the relevance of this mechanism to the pathophysiology of ALS, we examined *REST* and *UNC13A* expression with publicly available RNA-seq data for motor neurons isolated from the lumbar region of individuals with sporadic ALS by laser capture microdissection (**Figure 7A**). The symptoms of these individuals were first apparent rostrally and progressed in a descending manner, indicating that the motor neurons in the lumbar anterior horn were relatively preserved (Krach *et al*, 2018). We found that *REST* expression was increased in the spinal motor neurons of these samples compared with those of control individuals without ALS (**Figure 7B**).

**Figure 7.**
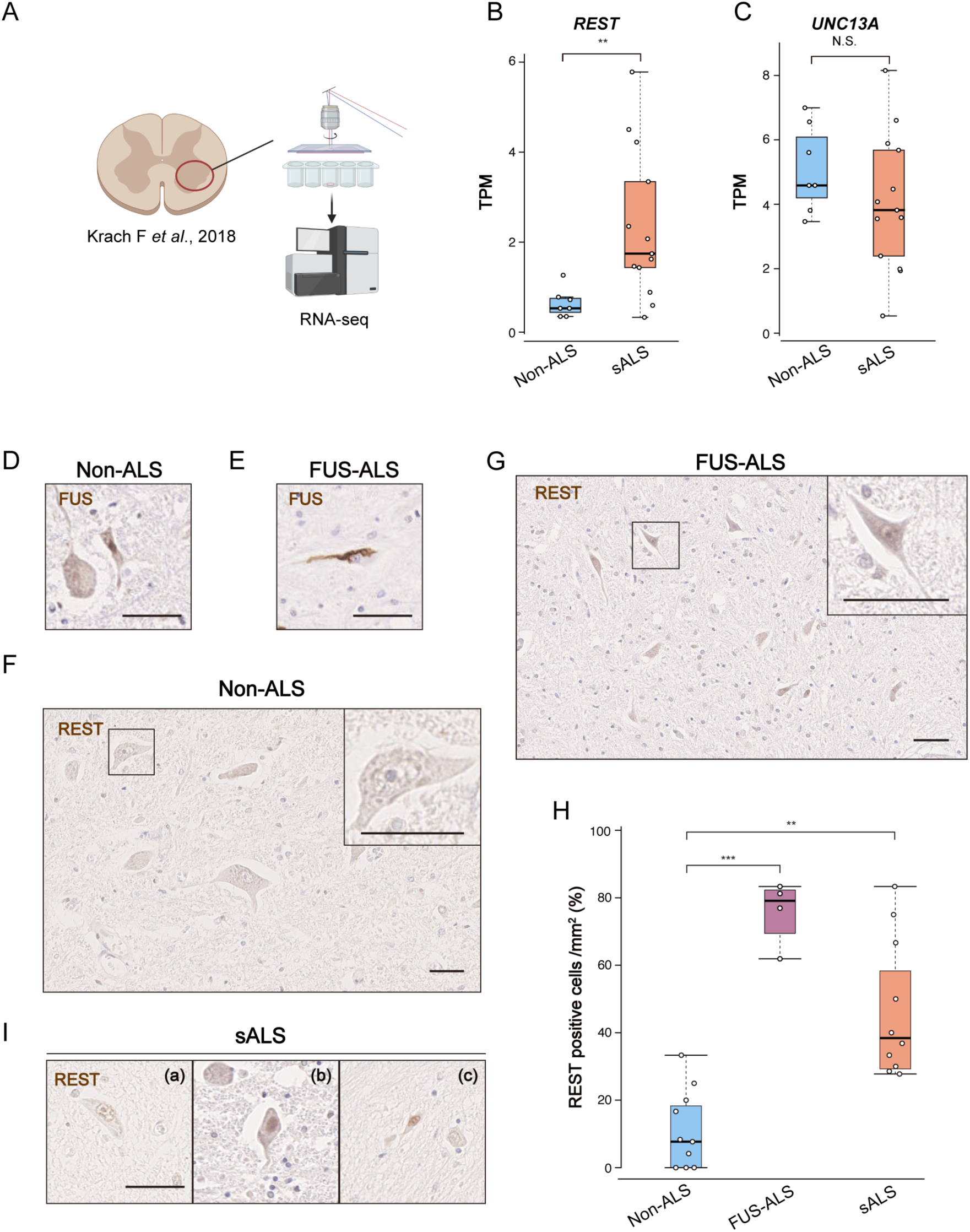
REST is overexpressed in spinal motor neurons of individuals with ALS. (A) RNA-seq analysis was previously performed for lumbar motor neurons isolated from control (non-ALS) individuals (*n* = 7) and patients with sporadic ALS (sALS, *n* = 13) by laser capture microdissection (Krach *et al*., 2018). The data are available under accession number GSE76220 in the GEO database. The illustration was created with Biorender.com. (B and C) *REST* (B) and *UNC13A* (C) expression for the samples in (A). TPM, transcripts per million. Data are presented as box plots, in which the boxes show the median and upper and lower quartile values, and the whiskers represent the range. \*\**p* < 0.01, NS (Mann-Whitney U test). (D and E) Immunohistochemical staining for FUS in spinal motor neurons of a control (non-ALS) individual with sporadic inclusion body myositis (sIBM) and an individual with familial ALS associated with a *FUS* mutation (R521C/+), respectively. Scale bars, 50 μm. (F and G) Immunohistochemical staining for REST in spinal motor neurons of a control individual with sIBM and an individual with familial ALS associated with a *FUS* mutation (R521C/+), respectively. The boxed regions in the main images are shown at higher magnification in the insets. Scale bars, 50 μm. (H) Quantification of REST-positive cells among anterior horn neurons by immunohistochemical analysis for three control individuals (with sIBM, carcinoma peritonitis, or multiple system atrophy), one individual with familial ALS associated with a *FUS* mutation (R521C/+), and three individuals with sALS. The proportion of REST-positive cells was quantified per square millimeter in four sections per sample. \*\**p* < 0.01, \*\*\**p* < 0.001 (Kruskal-Wallis test followed by Dunn’s test). See also **Figure S6C** for the values for each individual. (I) Immunohistochemical staining for REST in spinal motor neurons of three individuals with sALS. TDP-43 pathology was apparent in patients (b) and (c), but not in patient (a), as is shown in **Figure S6D**. Scale bar, 50 µm. See also **Figure S6**.

The attenuation of *STMN2* expression, previously identified as a potential pathological indicator of TDP-43 mislocalization (to the cytoplasm rather than the nucleus) (Prudencio *et al*, 2020), was apparent in the ALS samples of the RNA-seq data set (**Figure S6A**), suggestive of TDP-43 pathology. However, no correlation was detected between *STMN2* and *REST* expression levels (**Figure S6B**), suggesting that *REST* overexpression was independent of TDP-43 pathology. The expression level of *UNC13A* tended to be decreased in the individuals with sporadic ALS, but this difference did not achieve statistical significance (**Figure 7C**). Although the RNA-seq protocol specifically targeted motor neurons, the samples likely also included other cell types such as astrocytes and microglia (Krach *et al*., 2018). Given that *UNC13A* expression is minimal in glial cells (Uhlén *et al*, 2015), such contamination might have masked a potentially significant difference in *UNC13A* mRNA levels between non-ALS and ALS motor neurons.

We also examined REST protein expression in spinal motor neurons of individuals with ALS by immunohistochemical staining. In control individuals, FUS was localized predominantly to the nucleus (**Figure 7D**). In contrast, in an individual with ALS associated with a mutation in the NLS of FUS (FUS-ALS), FUS was localized mostly to the cytoplasm, where it formed aggregates (**Figure 7E**). Of note, the number of motor neurons expressing REST was significantly increased in FUS-ALS (**Figures 7F–7H**), suggestive of a pathological change linked to FUS dysfunction and involving REST overexpression. We also observed an increase in the number of REST-positive motor neurons in individuals with sporadic ALS compared with control individuals (**Figures 7H, 7I**, and **S6C**). Importantly, REST overexpression was observed even in an individual with sporadic ALS lacking TDP-43 pathology (**Figures 7I** and **S6D**), suggesting that REST overexpression is independent of such pathology. Together, these findings thus indicated that REST overexpression is a common pathological feature in both FUS-ALS and sporadic ALS, and they suggest a broad role for REST in ALS pathogenesis that is likely mediated by effects on the expression of synapse-related genes including *UNC13A*.

## Discussion

Multiple RBPs are closely associated with ALS both genetically and pathologically. However, a unified downstream pathway by which these RBPs contribute to ALS pathogenesis has remained to be identified. We have now investigated *UNC13A* as a potential convergence point for mechanisms by which the loss of function of four ALS-associated RBPs gives rise to ALS. UNC13A plays a pivotal role in the packaging of neurotransmitters into synaptic vesicles and the subsequent transport of these vesicles to the presynaptic membrane (Betz *et al*, 2001). In addition, it facilitates the priming and docking of the vesicles, ensuring that they are properly positioned for rapid neurotransmitter release on neuronal activation (Augustin *et al*., 1999; Siksou *et al*, 2009). UNC13A therefore maintains effective synaptic transmission and overall neural function. The prevalence of intronic mutations in *UNC13A* that destabilize the mRNA in individuals with ALS suggests that a loss of the UNC13A protein can contribute to ALS pathogenesis. Furthermore, restoration of UNC13A expression in TDP-43–depleted neurons derived from induced pluripotent stem cells (iPSCs) was found to fully rescue impaired presynaptic function (Keuss *et al*, 2024), suggesting that the downregulation of UNC13A that results from TDP-43 loss underlies the disruption of synaptic integrity apparent in ALS. Our findings now extend this observation by showing that functional impairment of additional ALS-associated RBPs also results in suppression of *UNC13A* expression, suggesting that *UNC13A* is a convergence point for mechanisms underlying the disruption of presynaptic vesicle function. This pathophysiological convergence is consistent with evidence that synaptic dysfunction is a fundamental characteristic of ALS and is closely linked to the functional abnormalities of several ALS-related genes (Clayton *et al*, 2024). In the widely studied mouse model of ALS based on transgenic expression of the G93A mutant of superoxide dismutase 1 (SOD1), the fusion of synaptic vesicles at the neuromuscular junction has been found to be impaired (Vinsant *et al*., 2013). Furthermore, synaptic loss has been detected in presymptomatic carriers of a mutation in *C9orf72*, which is the gene most strongly associated with familial ALS and frontotemporal dementia (Malpetti *et al*, 2021). Similarly, ultrastructural alterations of synaptic vesicles are evident at presynaptic terminals of the motor cortex in heterozygous *Fus* mutant mice (Scekic-Zahirovic *et al*, 2021). It is therefore plausible that synaptic terminal abnormalities can trigger pathogenesis in sporadic ALS.

REST is highly expressed in embryonic and neural stem cells, and it plays a key role in neuronal differentiation by repressing neuron-specific genes (Andrés *et al*., 1999; Johnson *et al*., 2008; Lunyak & Rosenfeld, 2005). Although it is expressed primarily in nonneuronal versus neuronal cells, REST has been implicated in neurodegenerative diseases. In Alzheimer’s disease, the loss of REST from the nucleus of neurons is associated with a loss of its neuroprotective function and contributes to cognitive impairment (Lu *et al*, 2014). Conversely, in Huntington’s disease, the mutated huntingtin protein induces abnormal REST accumulation in the nucleus and consequent suppression of the expression of genes such as that for brain-derived neurotrophic factor (BDNF) that are required for neuronal survival, thereby contributing to neurodegeneration (Zuccato *et al*, 2007; Zuccato *et al*, 2003). Although the possibility of a direct link between REST overexpression and ALS has not been well explored, studies have suggested that such a link may exist. In a mouse model of spinal cord injury, for example, REST disrupts axon regeneration by repressing regeneration-associated genes (Cheng *et al*, 2022). Furthermore, neurite regrowth following axotomy was found to be impaired in motor neurons derived from iPSCs harboring an ALS-associated *FUS* mutation (Stoklund Dittlau *et al*, 2021), suggesting the possibility that FUS dysfunction might lead to the overexpression or activation of REST, which in turn might be responsible for the inhibition of neuronal regeneration. In addition, REST overexpression in mice gives rise to impairment of spontaneous locomotion (Lu *et al*, 2018), a phenotype also seen in the SOD1(G93A) mouse model of ALS (Allodi *et al*, 2021), supporting the hypothesis that REST plays a role in ALS pathogenesis.

We have now shown that MATR3, FUS, and hnRNPA1 bind to *REST* mRNA and thereby attenuate *REST* expression at the posttranscriptional level. This finding is consistent with previous RNA-seq data showing that *REST* mRNA stability increased in response to FUS depletion in human neural progenitor cells in which RNA synthesis had been inhibited (Kapeli *et al*, 2016). Of note, we further show that the 27YS mutant of FUS is deficient in liquid-liquid phase separation (LLPS) and unable to downregulate *REST* mRNA in FUS-KO cells, underscoring the importance of such phase separation in such regulation by FUS. Both FUS and hnRNPA1 harbor large intrinsically disordered PrLDs that facilitate LLPS, with many ALS-associated mutations having been found to concentrate in these regions (Milicevic *et al*, 2022). Although MATR3 lacks a distinct PrLD, its NH_2_-terminal region contains a disordered domain that mediates LLPS, and the S85C mutation of MATR3, which is strongly linked to ALS onset, is located in this region and influences the condensation process (Johnson *et al*., 2014). In addition to the RBPs studied here, others including ATXN2, TIA-1, hnRNPA2/B1, and TAF15 have been found to harbor ALS-related mutations in PrLDs (Mann & Donnelly, 2021; Milicevic *et al*., 2022). The structural commonalities among these ALS-associated RBPs suggest that they may share a common pathophysiological mechanism. We hypothesize that the LLPS potential shared by FUS, hnRNPA1, MATR3, and possibly other ALS-associated RBPs with disordered domains may play a key role in *REST* mRNA downregulation.

We found that REST is overexpressed in motor neurons of individuals with sporadic ALS. A recent study showed that dysfunction of TDP-43 in sporadic ALS induces dysregulation of target RNA metabolism while the protein is still localized within the nucleus, before its mislocalization to the cytoplasm (Rothstein *et al*, 2023). Whereas mislocalization of RBPs other than TDP-43 is not frequently encountered in sporadic ALS (Honda *et al*., 2015; Tada *et al*., 2018; Tyzack *et al*., 2019), similar dysfunction of FUS, MATR3, and hnRNPA1 might also be evident while they remain in the nucleus. We speculate that such impaired function of these ALS-associated RBPs might lead to the upregulation of REST, potentially contributing to the repression of synapse-related genes such as *UNC13A*, in the motor neurons of individuals with sporadic ALS. With regard to the potential therapeutic approach of restoring UNC13A expression with a splice-correcting antisense oligonucleotide in individuals with ALS associated with TDP-43 pathology (Keuss *et al*., 2024), it may also be important to take into account the potential overexpression of REST in motor neurons of such patients.

In summary, we have revealed that the loss of each of four RBPs may contribute to ALS pathogenesis by convergence on a common pathophysiological pathway initiated by the downregulation of UNC13A expression. The mechanism by which the loss of TDP-43 gives rise to the attenuation of UNC13A expression is distinct from that by which the loss of MATR3, FUS, or hnRNPA1 does so, with the latter mechanism being mediated by transcriptional repression of *UNC13A* in a manner dependent on the upregulation of REST. The identification of this mechanism may provide a basis for the development of new therapeutic agents for ALS that target REST activity.

## Methods

### Plasmid construction

For construction of Cas9-sgRNA plasmids, sgRNAs were designed with the use of CRISPR direct (https://crispr.dbcls.jp) and subcloned into the pSpCas9(BB)-2A-Puro (PX459) V2.0 vector (Addgene) (Ran *et al*, 2013). Lentivirus vectors for doxycycline-inducible NH_2_-terminally FLAG-tagged human FUS (WT and deletion mutants) were kindly provided by T. Nakaya.(Nakaya, 2020) For construction of the doxycycline-inducible 3×FLAG-hnRNPA1 vector, cDNA encoding human hnRNPA1 was initially cloned into the pENTR vector (Thermo Fisher Scientific) and verified by sequencing. The hnRNPA1 sequence of the resulting vector was then transferred by recombination with the use of LR Clonase II (Thermo Fisher Scientific) into a p3xFLAG vector that had been modified to include attR sites. The 3×FLAG-tagged hnRNPA1 cDNA was subsequently amplified from the p3xFLAG vector by PCR and cloned with the use of an In-Fusion Cloning Kit (Takara Bio) into the pEN_TTGmiRc2 vector (Addgene), from which the EGFP and miR-30a coding regions had previously been removed with restriction enzymes. The resulting pEN_TTG-3xFLAG-hnRNPA1 vector was finally subjected to recombination with the pSLIK-neo destination plasmid (Addgene). Human MATR3 cDNA was cloned into the pENTR vector and subjected to similar verification and recombination steps as for hnRNPA1 cDNA, yielding a recombined PB-TA-ERN vector (Addgene) with the use of LR Clonase II. In addition, doxycycline-inducible lentivirus vectors for LLPS-deficient FUS mutants were constructed from cDNAs synthesized by GeneArt Custom Gene Synthesis (Invitrogen) and assembled in a similar manner to that adopted for the doxycycline-inducible 3×FLAG-hnRNPA1 construct. For construction of the WT luciferase vector for the *UNC13A* promoter, the promoter region of human *UNC13A* as defined in the UCSC genome browser was amplified from genomic DNA by PCR and then inserted into the pGL4.12 vector, which had previously been digested with BamHI and HindIII. A construct lacking the REST binding motif was generated with the use of a PrimeSTAR Mutagenesis Basal Kit (Takara Bio).

### Cell culture and transfection

HEK293T and SH-SY5Y cells were maintained in Dulbecco’s modified Eagle’s medium (DMEM) supplemented with 10% fetal bovine serum, penicillin (50 U/ml), streptomycin (50 μg/ml), 2 mM L-glutamine, 1% MEM–nonessential amino acids, and 1% sodium pyruvate. The cells were transiently transfected with plasmid DNA with the use of the PEI MAX reagent (Polyscience) and FuGENE HD (Promega), respectively.

### RNA-seq analysis

A TruSeq Standard mRNA LT Sample Prep Kit (Illumina) was used for library preparation. Sequencing was conducted on an Illumina HiSeq 2500 instrument to yield 151-nucleotide paired-end reads. Adapter sequences and low-quality bases were trimmed with the use of Trim_Galore. The resulting high-quality reads were aligned to the reference genome (GRCh38) with the use of STAR. Gene expression analysis, including k-means clustering and transcription factor motif analysis, was performed with iDEP (versions 2.01 and 96, respectively) (Ge *et al*, 2018).

### RNA interference

Cells were transfected twice with 20 nM siRNAs with the use of the RNAiMax reagent (Thermo Fisher Scientific). The second transfection was performed 72 h after the onset of the first, and the cells were harvested for analysis 24h after the second transfection.

### RT-PCR and RT-qPCR

RNA was isolated from cells with the use of an SV Total RNA Isolation System (Promega) and was subjected to RT with a PrimeScript RT Reagent Kit (Takara Bio). For qPCR, the resulting cDNA was amplified by real-time PCR analysis with the use of a StepOnePlus Real-Time PCR System (Life Technologies) and Fast SYBR Green Master Mix (Life Technologies). Data were analyzed with the 2-ΔΔCT method and normalized by the amount of human *GAPDH* mRNA. For PCR, cDNA was amplified with the use of PrimeSTAR Max DNA Polymerase (Takara Bio), and the amplification products were subjected to 2% agarose gel electrophoresis. The primer sequences for qPCR and PCR are listed in **Table S5**.

### Nascent RNA purification

Cells were seeded at a density of 1 × 10⁶ cells per well in six-well plates, cultured overnight, and labeled with 0.2 mM 4-EU for 1 h. Total RNA was extracted from the cells with the use of an SV Total RNA Isolation System (Promega), and nascent RNA was purified from the total RNA with a Click-iT Nascent RNA Capture Kit (Invitrogen). In brief, the extracted RNA was subjected to biotinylation by incubation with Click-iT Reaction Buffer, Biotin Azide, and CuSO₄ solution for 30 min at room temperature with vortex mixing. Biotinylated nascent RNA was captured with Dynabeads MyOne Streptavidin T1 magnetic beads (Invitrogen). The bead-bound RNA was washed and then immediately subjected to cDNA synthesis with a SuperScript VILO cDNA Synthesis Kit (Thermo Fisher).

### Immunoblot analysis

Cells were washed with phosphate-buffered saline and then lysed for 10 min at 4°C in NP-40 lysis buffer (0.5% Nonidet P-40, 50 mM Tris-HCl [pH 7.5], 150 mM NaCl, 10% glycerol) supplemented with a protease inhibitor cocktail (aprotinin [10 μg/ml, Sigma], leupeptin [10 μg/ml, Peptide Institute], 1 mM phenylmethylsulfonyl fluoride [Wako]). The lysates were centrifuged at 20,000 × g for 15 min at 4°C, and the resulting supernatants were harvested for immunoblot analysis. For preparation of an insoluble fraction, the pellet obtained by the centrifugation step was solubilized by ultrasonic treatment in urea buffer (7 M urea, 2 M thiourea, 1% CHAPS detergent, 30 mM Tris-HCl [pH 8.0], 25 mM imidazole). For immunoblot analysis, the samples were mixed with Laemmli buffer and fractionated by SDS-polyacrylamide gel electrophoresis The separated proteins were transferred to a polyvinylidene difluoride membrane (Millipore), which was then incubated consecutively with primary antibodies, horseradish peroxidase–conjugated secondary antibodies, and chemiluminescence reagents. Signals were detected with a ChemiDoc Touch System (Bio-Rad).

### Generation of RBP-KO cell lines

Cells were transiently transfected with Cas9-sgRNA plasmids and exposed to puromycin (5 μg/ml) for 2 days, and the surviving cells were cloned by the limiting dilution method. The single cell–derived clones were expanded and then validated by genomic PCR analysis of extracted DNA followed by sequencing. The target sequences of the sgRNAs were as follows: 5′-CCCATG GAAAACAACCGAAC-3′ within *TARDBP*, 5′-CCAGCAGTCATCTCTCAGTA-3′ within *MATR3*, 5′-CGGACATGGCCTCAAACGgt-3′ within *FUS*, and 5′-TGCCGTCATGTCTAAGTCAG-3′ within *HNRNPA1*.

### Restoration of RBP expression in RBP-KO cell lines

For doxycycline-inducible FLAG-FUS (WT and mutant) or 3×FLAG-hnRNPA1 expression in corresponding KO cell lines, the FUS-KO or hnRNPA1-KO cells were infected with lentiviruses in the presence of polybrene (8 μg/ml) and then subjected to selection with G418 (400 μg/ml, Wako) for at least 7 days. The lentiviruses were produced by transfection of HEK293T cells with the pSLIK-neo vectors containing the rtTA-TRE–regulated FLAG-FUS or 3×FLAG-hnRNPA1 cDNA sequences, as well as with the packaging plasmid psPAX2 and the envelope plasmid pMD2.G (Addgene). For doxycycline-inducible MATR3 expression in MATR3-KO cells, the cells were transfected with the corresponding PiggyBac vector and the Super PiggyBac Transposase Expression Vector (System Biosciences) and were then subjected to selection with G418 (400 μg/ml) for 2 weeks. For induction of each RBP, cells were treated with doxycycline (1 μg/ml, LKT Laboratories) for 2 days.

### Analysis of pre-mRNA

Total RNA was extracted from cells with the use of an SV Total RNA Isolation System (Promega) and was subjected to RT with a PrimeScript RT Reagent Kit (Takara Bio) but without an oligo(dT) primer. The RT reaction was also performed with (+RT) or without (– RT) reverse transcriptase in order to control for genomic DNA contamination. The reaction products were subjected to qPCR analysis with primers designed to target an intronic region of the *UNC13A* gene. The abundance of *UNC13A* pre-mRNA was compared across cell lines by normalization of the cycle threshold (CT) values obtained from the +RT samples by those from the –RT samples.

### Luciferase reporter assay

The human *UNC13A* promoter region spanning nucleotides –200 to +121 relative to the TSS was amplified by PCR and ligated into the pGL4.12 luciferase reporter vector (Promega) to yield pGL4.12-UNC13A-WT. The pGL4.12-UNC13A-ΔR vector for the deletion mutant lacking an intact REST binding site was constructed with a PrimeSTAR Mutagenesis Basal Kit (Takara Bio). Luciferase activities of cell lysates were measured with a Dual-Glo Luciferase Assay System (Promega) and a Berthold Centro LB960 instrument. The ratio of firefly to *Renilla* luciferase activity was calculated. For knockdown of REST in HEK293T cells, the cells were transfected with Cas9-sgRNA vectors the day before transfection with the promoter-luciferase constructs. The target sequences of the sgRNAs were as follows: sgRNA pair #1, 5′-GTTATGGCCACCCAGGTAAT-3′ and 5′-AGACATATGCGTACTCATTC-3′; sgRNA pair #2, 5′-CAACAGTGAGCGAGTATCAC-3′ and 5′-GTCTTCTGAGAACTTGAGTA-3′.

### RIP assay

RIP was performed with a RIP-Assay Kit (MBL). In brief, cells were lysed in RIP lysis buffer supplemented with 1.5 mM dithiothreitol, RNase OUT (50 U/ml, Invitrogen), and protease inhibitors. The lysates were incubated for 3 h at 4°C under gentle rotation with Pierce Protein G Plus Agarose (Thermo Fisher Scientific) conjugated to antibodies specific for FUS (Santa Cruz Biotechnology, sc-47711), MATR3 (Abcam, ab151739), or hnRNPA1 (Santa Cruz Biotechnology, sc-56700). Mouse IgG (Santa Cruz Biotechnology, sc-2025) and rabbit IgG (Sigma, I5006) were used as controls. The beads were then washed extensively to remove nonspecifically bound material, after which coprecipitated RNA was isolated from the beads with an SV Total RNA Isolation System (Promega). The isolated RNA was subjected to RT with a PrimeScript RT Reagent Kit (Takara Bio), and the resulting cDNA was subjected to PCR amplification followed by 2% agarose gel electrophoresis for detection of *REST* mRNA.

### ChIP-qPCR analysis

ChIP assays were performed with the use of a SimpleChIP Enzymatic Chromatin IP Kit (Cell Signaling). In brief, cells (∼1 × 10^7^) were fixed with 1% formaldehyde for 10 min at room temperature, subjected to quenching, and enzymatically digested for 20 min at 37°C. Chromatin was sheared by ultrasonic treatment (five 30-s applications) and incubated overnight at 4°C with antibodies to REST (Millipore). Chromatin in immune complexes was then precipitated by incubation with protein G–conjugated magnetic beads, washed, and eluted. After reversal of cross-links and purification, precipitated DNA was subjected to qPCR analysis with specific primers (**Table S5**).

### Immunohistochemistry

Patients were diagnosed with ALS according to the revised El Escorial criteria (Brooks *et al*, 2000), with diagnoses being further confirmed pathologically by postmortem examination. Tissue from each level of the spinal cord was either immediately placed in 10% buffered formalin and embedded in paraffin for neuropathologic examination. Immunohistochemistry for FUS and TDP-43 was performed as previously described.(Akiyama *et al*., 2019) For REST staining, tissue sections were depleted of paraffin with xylene, rehydrated with a graded series of ethanol solutions in phosphate-buffered saline, and subjected to antigen retrieval by microwave irradiation for 20 min in EDTA buffer (pH 9.0). Endogenous peroxidase activity was blocked by treatment with 0.3% H_2_O_2_ in methanol for 30 min, after which the sections were exposed to Protein Block (Genostaff), processed with an Avidin/Biotin Blocking Kit (Vector), and incubated consecutively overnight at 4℃ with rabbit monoclonal antibodies to REST (Bethyl), for 30 min at room temperature with biotin-conjugated goat antibodies to rabbit IgG (Vector), and for 5 min at room temperature with peroxidase-conjugated streptavidin (Nichirei). Peroxidase activity was visualized by staining with diaminobenzidine, and the sections were counterstained with Mayer’s hematoxylin (Muto), dehydrated, and mounted with Malinol (Muto).

### Quantification and statistical analysis

Relative band intensities were quantified by densitometry with the use of ImageJ.(Schneider *et al*, 2012) Statistical analysis was performed with R software. Data were compared between two groups with the unpaired two-tailed Student’s *t* test or Mann-Whitney U test, and among three or more groups either by one-way analysis of variance (ANOVA) followed by Tukey’s post hoc test, or by the Kruskal-Wallis test followed by Dunn’s test. Data are presented as means ± SEM unless indicated otherwise, and a *p* value of <0.05 was considered statistically significant.

## Acknowledgments

This work was supported by KAKENHI grants (22K15702 to Y.W., 21K07411 to N.S., 23H02821 to M.A., 21H02458, 24K02300 to K.N.) from the Japan Society for the Promotion of Science (JSPS), by the Japan ALS Association (JALSA, to Y.W.), and by the Yukihiko Miyata Memorial Trust for ALS Research (to Y.W.). We thank Y. Nagasawa and M. Kikuchi for general technical and secretarial assistance; H. Aoyama for assistance with immunohistochemistry; other laboratory members for discussion; and T. Nakaya (International University of Health and Welfare) for providing pSLIK-neo-FLAG-FUS and corresponding mutant plasmids.

## Author contributions

Y.W. conceptualized and designed the research, conducted the experiments, and wrote the manuscript. K.N. directed the study and cowrote the manuscript. N.S. supervised the study and contributed to immunohistochemistry. T.N., M.H., and T.A. reviewed the results and provided guidance. H.W. and M.A. provided advice. All authors discussed the results and commented on the manuscript.

## Disclosure and competing interests statement

The authors declare no competing interests.

## Supplementary Figures

**Figure S1.**
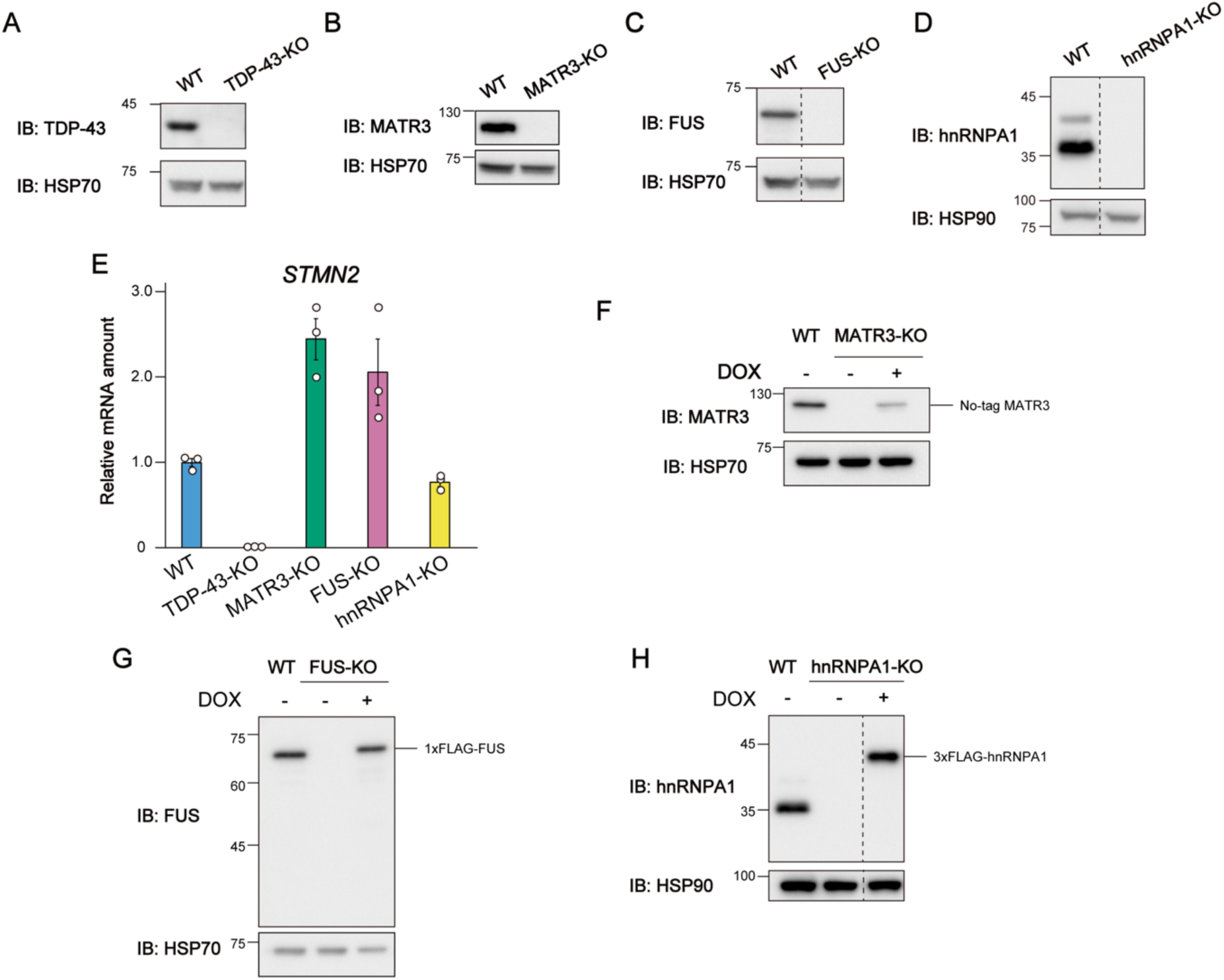
Generation of RBP-KO cell lines and rescue of corresponding RBP expression, related to Figure 1. (A–D) Immunoblot (IB) analysis of RBPs in WT and TDP-43-KO (A), MATR3-KO (B), FUS-KO (C), and hnRNPA1-KO (D) cells. HSP70 or HSP90 served as a loading control. (E) RT-qPCR analysis of *STMN2* mRNA in WT and RBP-KO cell lines. Data are means ± SEM from three independent experiments. (F–H) Immunoblot analysis of RBPs in WT cells and in MATR3-KO (F), FUS-KO (G), and hnRNPA1-KO (H) cells complemented with a corresponding doxycycline-inducible RBP vector and exposed (or not) to doxycycline (DOX).

**Figure S2.**
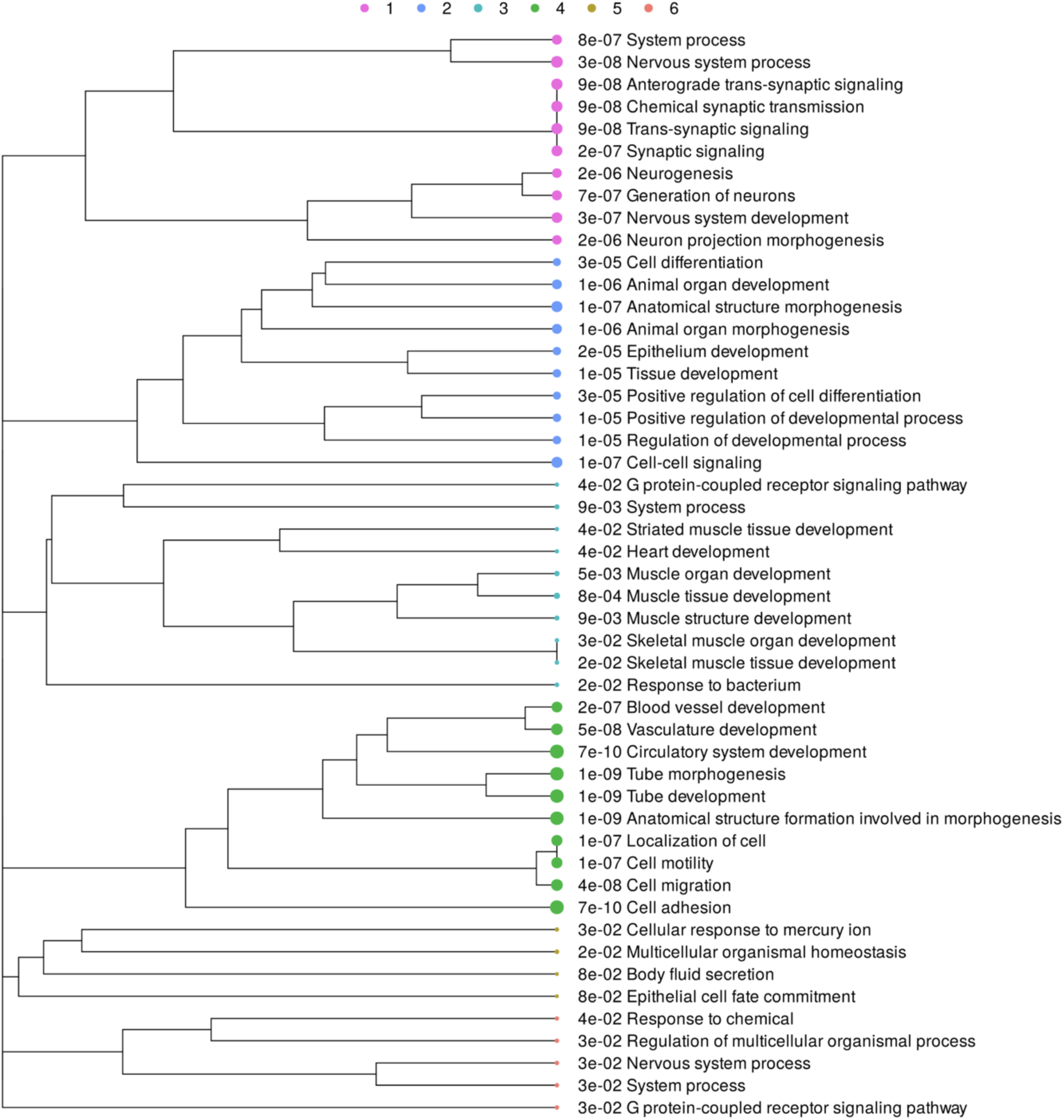
GO analysis of RNA-seq data from RBP-KO cells, related to Figure 1. Genes in clusters 1 to 6 shown in Figure 1B were subjected to GO biological process analysis. The numbers indicate the FDR values of each GO term, and the circle size indicates fold enrichment.

**Figure S3.**
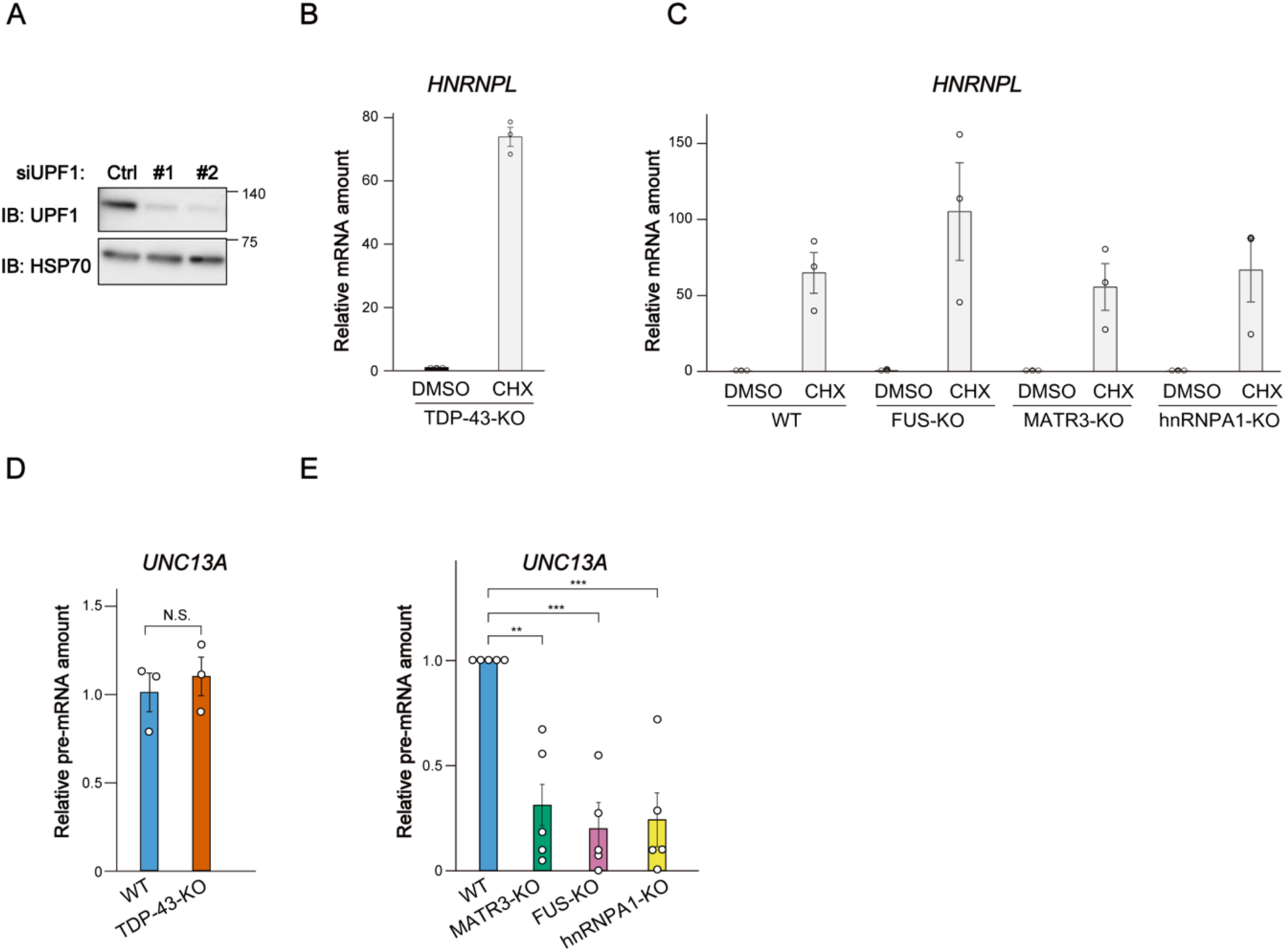
Inhibition of NMD and detection of *UNC13A* pre-mRNA in RBP-KO cells, related to Figure 2. (A) Immunoblot analysis of UPF1 in TDP-43-KO cells transfected with a GC duplex (negative control) or either of two siRNAs targeting *UPF1*. (B and C) RT-qPCR analysis of *HNRNPL* mRNA (positive control sensitive to NMD) in TDP-43-KO cells (B) or in WT, MATR3-KO, FUS-KO, and hnRNPA1-KO cells (C) after treatment with cycloheximide or DMSO vehicle. Data are means ± SEM from three independent experiments. (D) RT-qPCR analysis of *UNC13A* pre-mRNA in WT and TDP-43-KO cells. Data are means ± SEM from three independent experiments. N.S. (Student’s *t* test). (E) RT-qPCR analysis of *UNC13A* pre-mRNA in WT, MATR3-KO, FUS-KO, and hnRNPA1-KO cells. Data are means ± SEM from five independent experiments. \*\**p* < 0.01, \*\*\**p* < 0.001 (one-way ANOVA followed by Tukey’s post hoc test).

**Figure S4.**
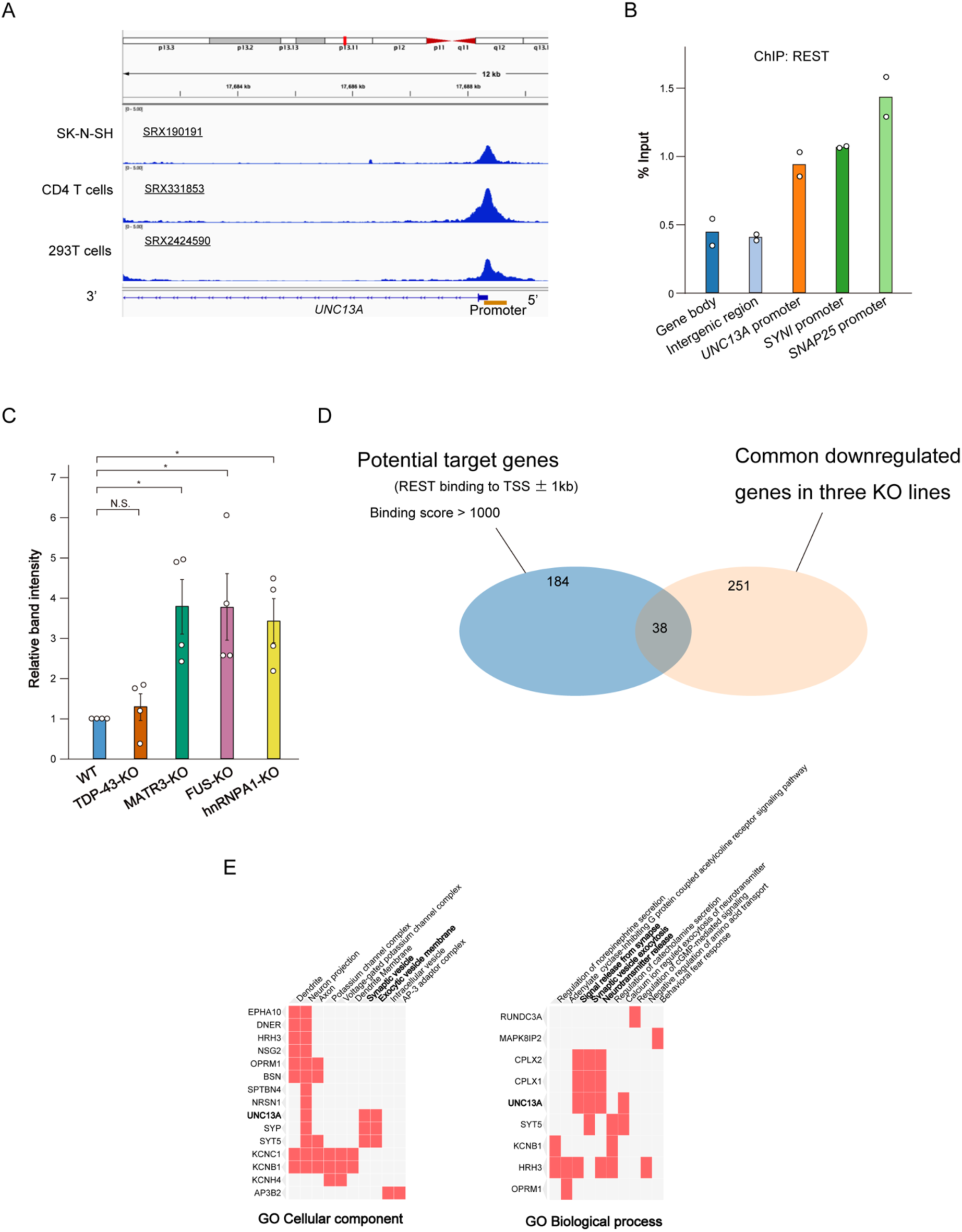
REST binds to the *UNC13A* promoter and suppresses the expression of various genes in RBP-KO cells, related to Figure 3. (A) Alignment of ChIP-seq reads for REST to the *UNC13A* promoter with the use of ENCODE data sets. ChIP-seq data from SK-N-SH cells, HEK293T cells, and CD4 T cells show specific binding of REST to the *UNC13A* promoter region (chr19:17688234-17688572 in Hg38). (B) ChIP-qPCR analysis of REST binding to the *UNC13A* promoter in SH-SY5Y cells. The gene body of *UNC13A* and an intergenic region were examined as negative controls to which REST does not bind. The promoter regions of *SNAP25* and *SYN1*, both of which are known to bind REST, were examined as positive controls. Data are means from two independent experiments. (C) Quantification of the band intensity for REST (CST antibody preparation) normalized by that for HSP70 in immunoblots similar to that shown in Figure 3F. Data are means ± SEM for four independent experiments. \**p* < 0.05, NS (one-way ANOVA followed by Tukey’s post hoc test). (D) Venn diagram showing the overlap between potential target genes of REST and commonly downregulated genes in MATR3-KO, FUS-KO, and hnRNPA1-KO cells. Potential target genes, defined as those to which REST binds within ±1 kb of the TSS, were identified by cross-referencing ENCODE ChIP-seq data with ChIP-Atlas. Genes with a binding score as calculated with MACS2 and STRING of >1000 were considered potential target genes. (E) GO analysis of the 38 potential REST target genes identified among the commonly downregulated genes in the three RBP-KO cell lines as in (D). These genes were annotated for GO terms with the Enrichr tool (https://maayanlab.cloud/Enrichr).

**Figure S5.**
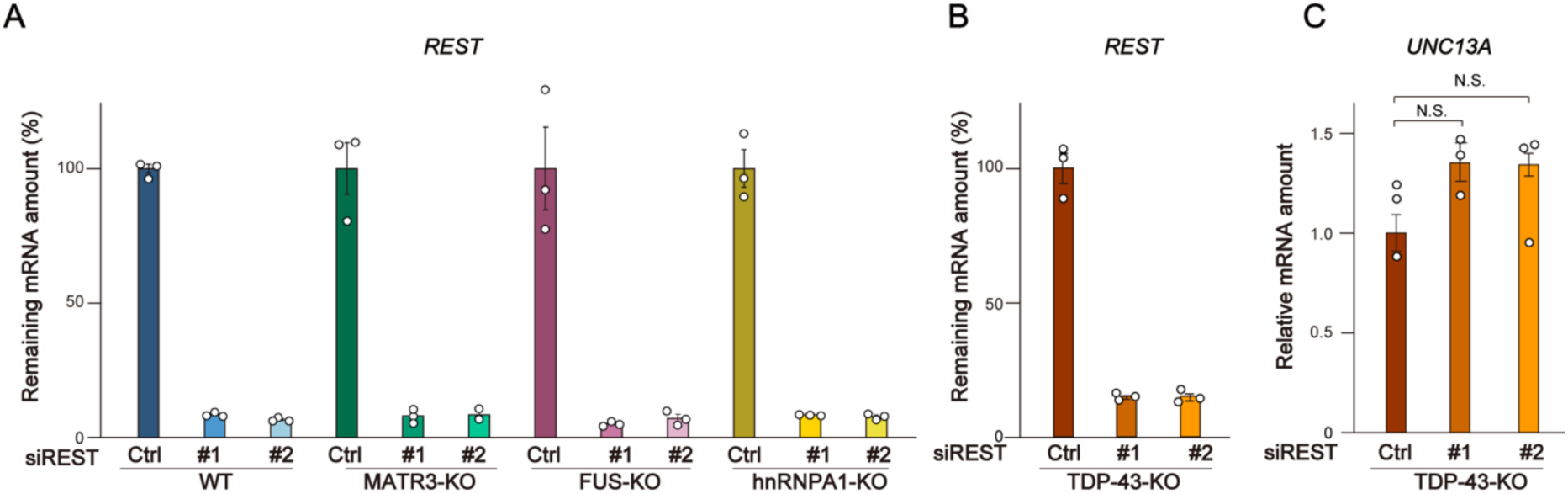
Knockdown of REST in RBP-KO cells, related to Figure 4. (A) RT-qPCR analysis of *REST* mRNA in WT, MATR3-KO, FUS-KO, and hnRNPA1-KO cell lines transfected with either a GC duplex (negative control) or one of two different REST siRNAs. Data are means ± SEM from three independent experiments. (B and C) RT-qPCR analysis of *REST* (B) and *UNC13A* (C) mRNAs in TDP-43-KO cells transfected as in (A). Data are means ± SEM from three independent experiments. N.S. (one-way ANOVA followed by Tukey’s post hoc test.)

**Figure S6.**
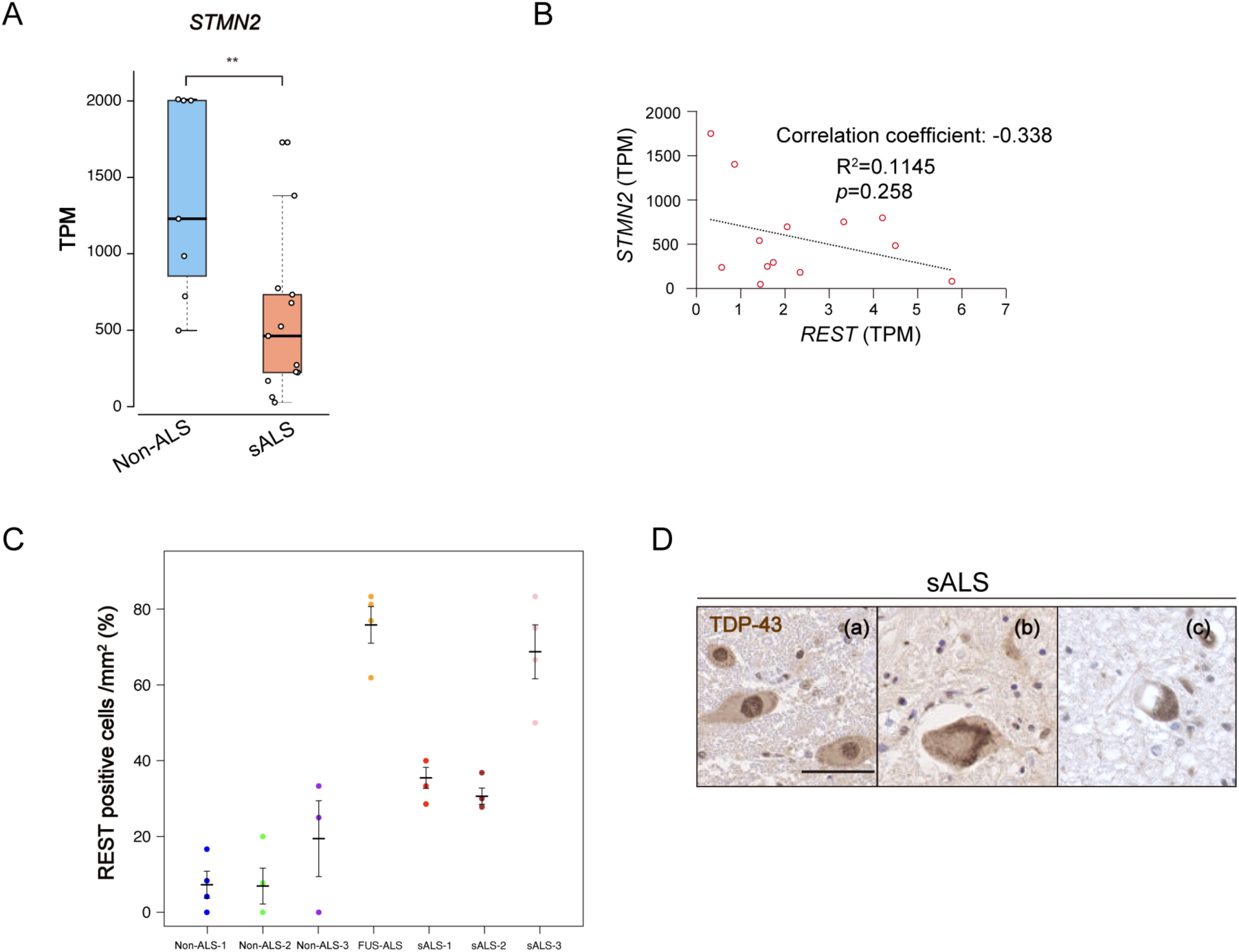
Overexpression of REST in individuals with familial or sporadic ALS, related to Figure 7. (A) Comparison of *STMN2* expression in lumbar motor neurons between control (non-ALS) individuals and individuals with sporadic ALS (sALS) in the GSE76220 data set. \*\**p* < 0.01 (Mann-Whitney U test). (B) Scatter plot showing the relation between the expression levels of *REST* and *STMN2* in lumbar motor neurons for sALS patients in the GSE76220 data set. Each red dot represents one sample. The dashed line indicates the linear regression fit. The R² value shows the proportion of variance in *STMN2* expression explained by *REST* expression, and the *p* value was calculated with Pearson’s correlation test. (C) Quantification of REST-positive cells among anterior horn neurons for three control individuals, an individual with familial ALS associated with a *FUS* mutation (R521C/+), and three sALS patients shown in Figure 7H. The percentage of REST-positive cells was quantified per square millimeter in four sections per sample. Both individual data points (four per sample) and their average are presented. (D) Immunohistochemical staining of TDP-43 in spinal motor neurons of three sALS patients. TDP-43 pathology was apparent in patients (b) and (c), but not in patient (a). Scale bar, 50 µm.

**Supplementary File**. Excel file containing Tables S1-S5

Table S1. The results of k-means clustering of RNA-seq

Table S2. ChIP-seq datasets for TFs binding to the *UNC13A* promoter region

Table S3. TF binding motif analysis in promoter regions of commonly downregulated genes Table S4. Predicted REST-binding promoters of commonly downregulated genes

Table S5. Primers used in this study

## Notes

### Competing Interest Statement

The authors have declared no competing interest.

